# The Mediator co-activator complex regulates Ty1 retromobility by controlling the balance between Ty1i and Ty1 promoters

**DOI:** 10.1101/176248

**Authors:** Alicia C. Salinero, Elisabeth R. Knoll, Z. Iris Zhu, David Landsman, M. Joan Curcio, Randall H. Morse

**Author notes:** Co-corresponding authors (MJC) and (RHM). MJC and RHM are Joint Senior Authors.

## Abstract

The Ty1 retrotransposons present in the genome of *Saccharomyces cerevisiae* belong to the large class of mobile genetic elements that replicate via an RNA intermediary and constitute a significant portion of most eukaryotic genomes. The retromobility of Ty1 is regulated by numerous host factors, including several subunits of the Mediator transcriptional co-activator complex. In spite of its known function in the nucleus, previous studies have implicated Mediator in the regulation of post-translational steps in Ty1 retromobility. To resolve this paradox, we systematically examined the effects of deleting nonessential Mediator subunits on the frequency of Ty1 retromobility and levels of retromobility intermediates. Our findings reveal that loss of distinct Mediator subunits alters Ty1 retromobility positively or negatively over a >10,000-fold range by regulating the ratio of an internal transcript, Ty1i, to the genomic Ty1 transcript. Ty1i RNA encodes a dominant negative inhibitor of Ty1 retromobility that blocks virus-like particle maturation and cDNA synthesis. These results resolve the conundrum of Mediator exerting sweeping control of Ty1 retromobility with only minor effects on the levels of Ty1 genomic RNA and the capsid protein, Gag. Since the majority of characterized intrinsic and extrinsic regulators of Ty1 retromobility alter a post-translational step(s), Mediator could play a central role in integrating signals that influence Ty1i expression to modulate retromobility.

**Author Summary:** Retrotransposons are mobile genetic elements that copy their RNA genomes into DNA and insert the DNA copies into the host genome. These elements contribute to genome instability, control of host gene expression and adaptation to changing environments. Retrotransposons depend on numerous host factors for their own propagation and control. The retrovirus-like retrotransposon, Ty1, in the yeast *Saccharomyces cerevisiae* has been an invaluable model for retrotransposon research, and hundreds of host factors that regulate Ty1 retrotransposition have been identified. Non-essential subunits of the Mediator transcriptional co-activator complex have been identified as one set of host factors implicated in Ty1 regulation. Here, we report a systematic investigation of the effects of loss of these non-essential subunits of Mediator on Ty1 retrotransposition. Our findings reveal a heretofore unknown mechanism by which Mediator influences the balance between transcription from two promoters in Ty1 to modulate expression of an autoinhibitory transcript known as Ty1i RNA. Our results provide new insights into host control of retrotransposon activity via promoter choice and elucidate a novel mechanism by which the Mediator co-activator governs this choice.

## Introduction

Retrotransposons have been extensively characterized as catalysts of evolutionary change and agents of genome instability [1-6]. In humans, retrotransposons have been shown to be upregulated in cancerous cells [7], and have been implicated in tumorigenesis [8]. Long terminal repeat (LTR) retrotransposons are of particular interest because they are the evolutionary progenitors of retroviruses [9], and have been found to be influenced heavily by host factors that control retroviral propagation [10-17].

Research focused on the LTR-retrotransposons in *Saccharomyces cerevisiae* has been fundamental to our understanding of the mechanism by which LTR-retrotransposons replicate, and how they interact with the host genome [2, 6, 18]. Ty1 is the most abundant and active of these LTR-retrotransposons, with 31 copies in the haploid genome of the common reference strain BY4741 and a mobility rate of approximately 1x10^-7^ to 1x10^-5^ per Ty1 element in each cell generation [19]. Retrotransposition initiates with transcription of the element (Fig 1A & B). The U3 region of the 5’ LTR contains promoter sequences recognized by DNA binding transcriptional activators, as well as a TATA box sequence (TATAAAAC) (Fig 1B). A second putative TATA box sequence at the 5’ LTR, with sequence TATTAACA, does not conform to identified functional TATA element sequences [18, 20].

**Fig 1:**
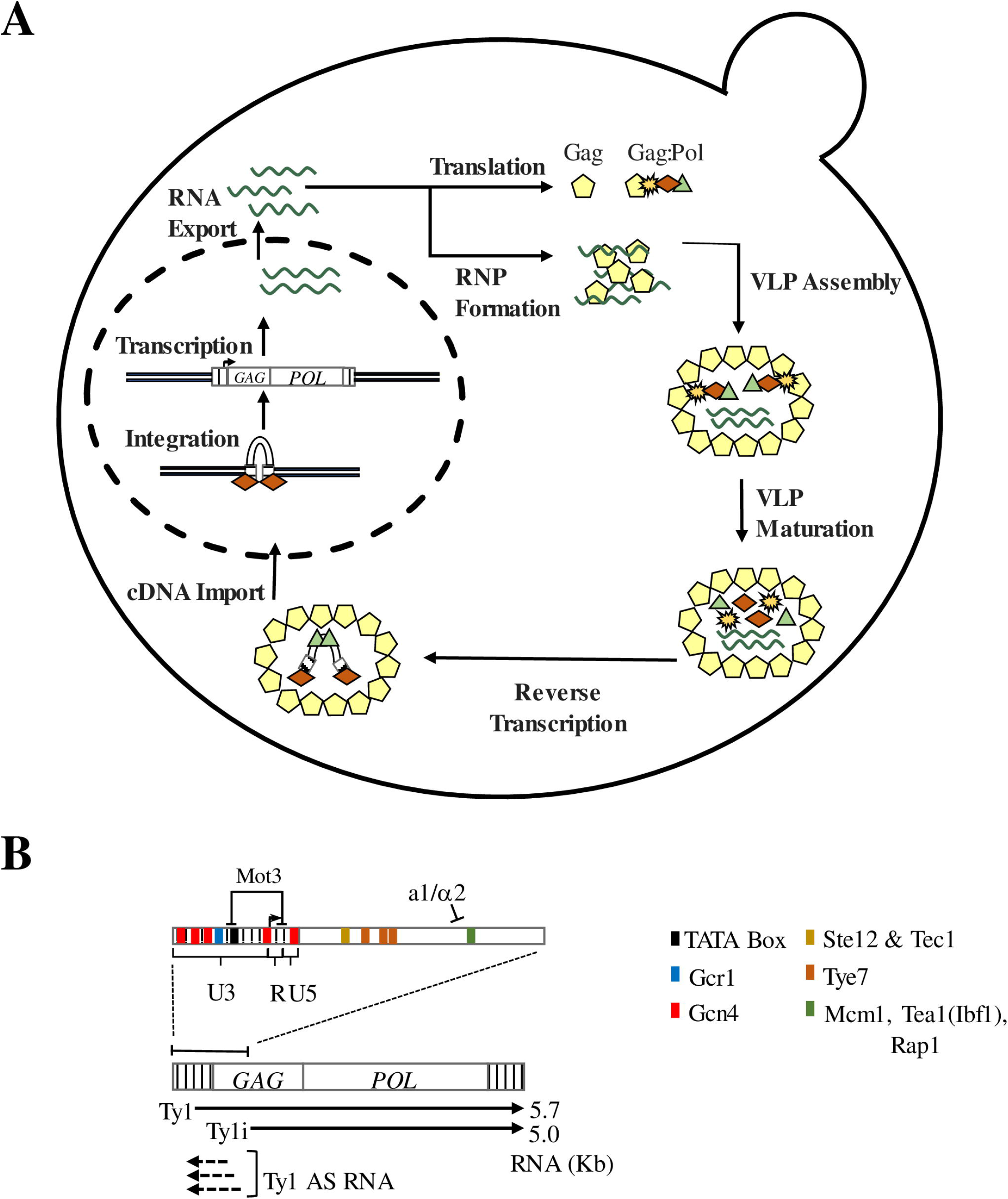
The Ty1 retrotransposon lifecycle and transcriptional regulation. (A) The Ty1 mobility lifecycle. Following transcription, Ty1 RNA is exported to the cytoplasm where it localizes co-translationally to a microscopically distinct cytoplasmic focus known as the retrosome. The retrosome is the site of assembly of the VLPs, which serve as the sites for Ty1 protein maturation and reverse transcription of Ty1 cDNA. Ty1 cDNA is then transported back into the nucleus and integrated into the host genome. (B) Genetic structure of Ty1. Ty1 contains two ORFs (GAG and *POL*) flanked by a 5 and a 3 LTR. Ty is transcribed from the 5 untranslated region (R-U5) in the 5 LTR beginning at nt +238, and it terminates within the 3 LTR. In addition to the primary Ty1 transcript, the Ty1i transcript initiates within the *GAG* ORF at nt +1000 to produce a 5.0 kb truncated transcript. Antisense (AS) RNA is also transcribed from within the *GAG* ORF to the 5 LTR. The Ty1 5 LTR as well as the first 1 kb of the *GAG* ORF contain several transcription factor binding sites, as well as a TATA element. Sites for Ste12, Tec1, Tye7, Mcm1, Tea1, and Rap1 occur downstream of the Ty1 TSS. The transcription termination sites in the 3 LTR (TS1 and TS2) are located in the R-U5 region.

The 5.7kb Ty1 transcript contains two partially overlapping open reading frames, *GAG* and *POL*, the latter of which is translated only when a specific +1 frameshift event occurs in the overlapping region (Fig 1B). This frameshifting event enables production of two translational products, the p49-Gag protein and the p199-Gag-Pol polyprotein. Ty1 protease, a factor that is encoded in the *POL* ORF, processes p49-Gag to p45-Gag as an integral part of Ty1 protein maturation. The Gag-Pol polyprotein is processed to yield p45-Gag, protease, reverse transcriptase and integrase. In addition to its protein coding function, the Ty1 transcript serves as a template for reverse transcription of the element. Ty1 protein processing and reverse transcription of Ty1 RNA occur within a cytoplasmic capsid of Gag protein known as the virus-like particle (VLP) (Fig 1A) [18]. Following reverse transcription, Ty1 cDNA is transported back to the nucleus and integrated into the host genome through the activity of integrase (Fig 1A) [21, 22].

Ty1 elements also produce a second transcript that initiates at +1000 bp in the Ty1 element, 762 bp downstream of the Ty1 transcription start site (TSS) (Fig. 1B). This transcript, now termed Ty1i, is readily observed in *spt3* and *snf5* mutants in which transcript levels of full-length Ty1 are severely reduced, but is difficult to detect against the background of Ty1 in wild type cells [23-25]. Ty1i RNA is important in copy number control (CNC), which is defined by a copy number-dependent decrease in Ty1 retrotransposition observed both in *S. cerevisiae* and its close relative, *S. paradoxus* [24, 26, 27]. CNC is enforced by a dominant, *trans*-acting regulatory protein known as p22-Gag that is translated from Ty1i RNA [24, 28-31]. This truncated Gag protein, as well as its proteolytically processed form, p18-Gag, retains the ability to associate with p49-or p45-Gag. However, incorporation of p22-Gag into the VLP (where it is processed to p18-Gag) disrupts nucleocapsid formation, thereby halting Ty1 protein maturation and production of Ty1 cDNA [24, 28, 29, 32].

Ty1 relies extensively on autoregulatory factors and host factors to successfully complete its mobility cycle and limit its mobility so as not to destabilize the host genome [18, 33]. Host factors that regulate Ty1 mobility include subunits of the Mediator transcriptional co-activator complex [34-39]. Mediator plays a crucial role in the formation of the PIC at all Pol II transcribed genes, in part by acting as a bridge between DNA binding transcriptional activator proteins and the RNA Pol II transcription machinery [40-47]. In *Saccharomyces cerevisiae*, Mediator is a 1.4 MDa complex composed of 25 individual subunits organized into four modules (Fig 2A) [48-51]. The core Mediator complex contains the “head,” “middle,” and “tail” modules, while a fourth kinase module is transiently associated with the core complex in a context-specific manner [52]. The tail domain is generally responsible for Mediator’s association with transcriptional activator proteins, while the head and middle are involved in association of RNA Polymerase II (Pol II) and pre-initiation complex (PIC) formation [42, 50].

**Fig 2:**
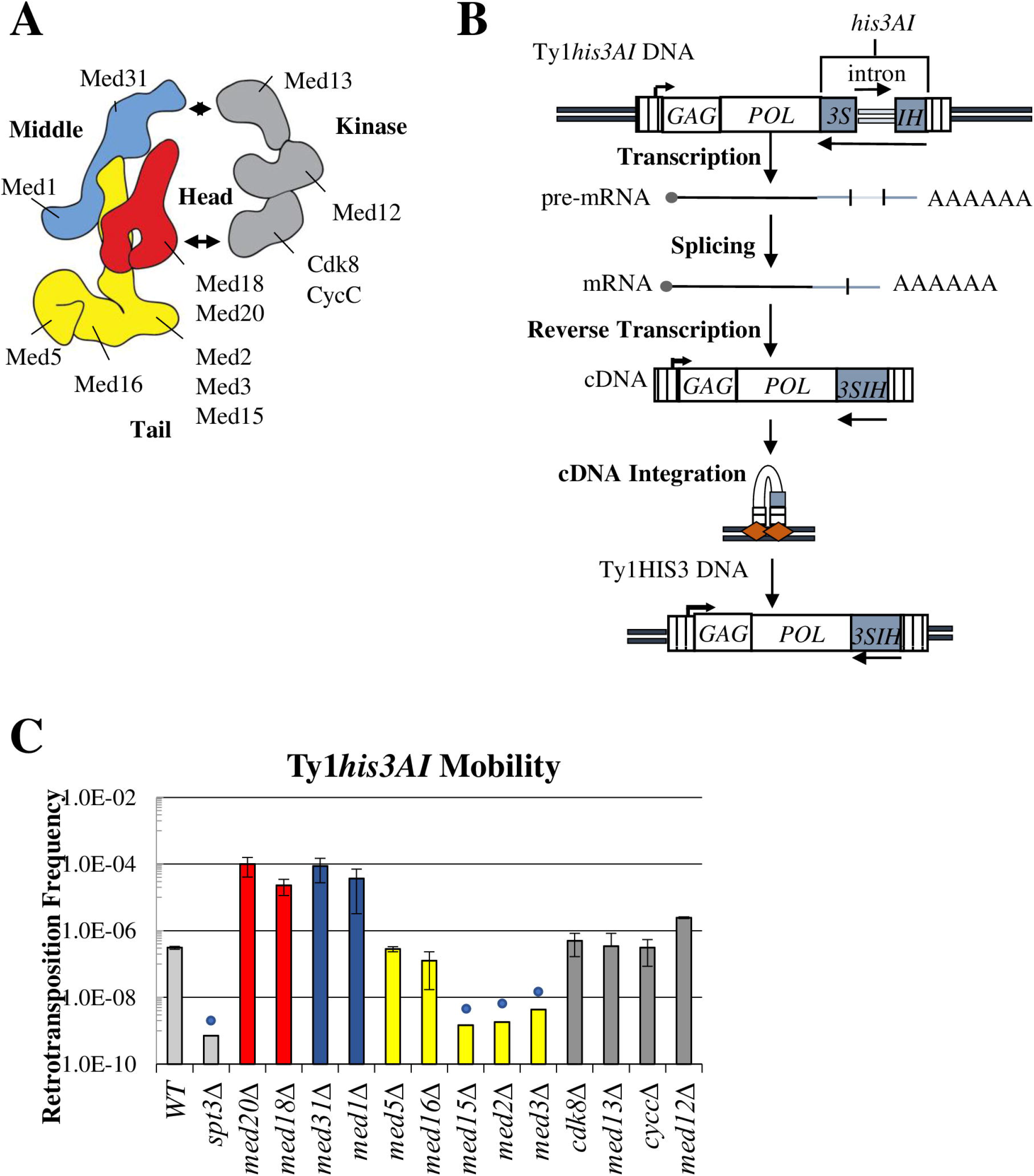
Mediator subunit deletions influence Ty1 mobility in a module-specific manner. (A) The Mediator transcriptional coactivator complex is composed of head (red), middle (blue), tail (yellow), and kinase (grey) modules (figure based on recent cryo-EM structure [50]). Individual subunits investigated in this study are labeled. (B) Schematic of the retromobility assay [19]. The *HIS3* ORF is inserted into *POL* in the opposite orientation of Ty1 and contains an intron that is in the sense orientation relative to Ty1. Splicing of this intron from the Ty1his3AI transcript, followed by integration, results in His+ colonies. (C) The frequency of retrotransposition, shown on a log scale, of the chromosomal Ty1his3AI-3114 element was measured in congenic WT, *spt3*Δ and Mediator subunit deletion strains. Error bars represent standard deviation for three biological replicates. Blue circles denote values that represent upper limit retrotransposition estimates in strains with no retrotransposition events among the total number of cells assayed. Bars are color-coded to match structural organization as shown in (A).

Several studies have implicated individual Mediator subunits as either activators or repressors of Ty1 retromobility. While no concerted effort to determine the role of all Mediator non-essential subunits on Ty1 mobility has been reported, individual studies have reported that loss of individual Mediator subunits affects a step in retrotransposition between transcription and integration while having minor effects on Ty1 transcript levels [18, 34-39]. This conclusion is strikingly incongruous with Mediator’s canonical role as a transcriptional regulator. To date, no mechanistic characterization of Mediator’s positive and negative influences on Ty1 mobility has been undertaken, and an explanation for its apparent post-transcriptional function in Ty1 mobility has been elusive.

In this study, we systematically determine the effects of deleting non-essential subunits of the Mediator complex on various steps in Ty1 retrotransposition, from Ty1 and Ty1i RNA expression to completion of the retrotransposition event. We show that deletion of Mediator complex subunits results in substantial, module-specific effects on the level of Ty1 retrotransposition. Consistent with previous findings, we find that Mediator subunit deletions have minimal effects on the levels of Ty1 RNA and Gag protein, but do result in substantial changes in the level of unintegrated cDNA that correspond to changes in the level of retrotransposition. We also report that deletion of individual subunits of the tail module triad, Med2-Med3-Med15, increases recruitment of Mediator and Pol II to a secondary promoter within the Ty1 *GAG* ORF. Use of this internal promoter results in expression of Ty1i RNA, whose translation product, p22-Gag, is a potent inhibitor of VLP formation and Ty1 cDNA synthesis. In contrast, loss of Mediator head module subunits Med18 or Med20 decreases Mediator association with the internal Ty1i promoter and results in increased Ty1 mobility. Thus, Mediator subunits control a post-transcriptional step in Ty1 mobility by modulating transcription of Ty1i RNA. Based on these observations, we propose a mechanism in which Mediator regulates Ty1 retromobility by controlling the balance between utilization of Ty1 and Ty1i promoters.

## Results

### The Core Mediator Complex Has Profound Effects on Ty1 Retromobility

Previous genetic screens determined that individual Mediator subunits influence Ty1 retromobility through post-transcriptional mechanisms; however, the screens employed different assays for retromobility, and differed in their identification of specific Mediator subunits contributing to Ty1 retromobility [34, 35, 37, 38, 53]. To systematically investigate the role of all non-essential subunits of Mediator in Ty1 retromobility, a collection of strains, each containing a deletion of a non-essential Mediator subunit, was generated from a BY4741 progenitor strain containing a chromosomal *his3AI*-marked Ty1 element (S1 Table). These strains were then subjected to an established quantitative retromobility assay in which cells that sustain a retromobility event are detected as His+ prototrophs (Fig 2B) [19]. A mutant lacking the SAGA complex component Spt3 was chosen as a negative control for Ty1 retromobility due to the well-characterized requirement for Spt3 in Ty1 transcription and mobility [25, 54].

Results from this assay indicate that Mediator influences *Ty1his3AI* retromobility in a profound, module-specific manner (Fig 2C). Deletion of genes encoding subunits in the head or middle module increased Ty1 *his3AI* retromobility approximately 100-fold. Conversely, deleting any subunit from the Med2-Med3-Med15 tail module triad resulted in a retromobility level that was less than 1% of that of the wild-type strain, and below detection limits. The lack of Ty1his3AI retromobility in these mutants is not due to the loss of *HIS3* expression, since His+ prototrophs form readily when Ty1his3AI is driven by a heterologous promoter in *med3Δ* and *med15Δ* mutants, as demonstrated below. The Med2-Med3-Med15 subunits are direct targets of DNA-binding activator proteins [42, 47, 55, 56], and deletions of these subunits exhibit similar phenotypes [56-58]. In contrast, two other tail module subunits that exhibit distinct phenotypes when deleted, Med5 and Med16, do not appear to affect Ty1 retromobility. The transiently associated kinase module also does not substantially influence Ty1 retromobility. Kinase module deletion strains were omitted from further analysis in this work. The disparate effects of deleting subunits in different modules is consistent with previous data indicating that the Mediator complex regulates gene expression in a module-dependent manner [58].

### Mediator Influences Ty1 cDNA without Altering Ty1 Transcript or Gag Protein Levels

Given Mediator’s role as a transcriptional co-activator, we first sought to determine whether the changes in Ty1 mobility observed in the Mediator subunit deletion strains were caused by altered Ty1 transcript levels. Changes in Ty1 RNA levels in head, middle or tail module gene deletion strains relative to the wild-type strain, visualized by northern blot analysis, were modest and not statistically significant (*p*>0.1, one-way ANOVA) (Fig 3A). These data indicate that Mediator head, middle and tail subunits do not regulate Ty1 retromobility by affecting the steady-state level of genomic Ty1 transcripts.

**Fig 3:**
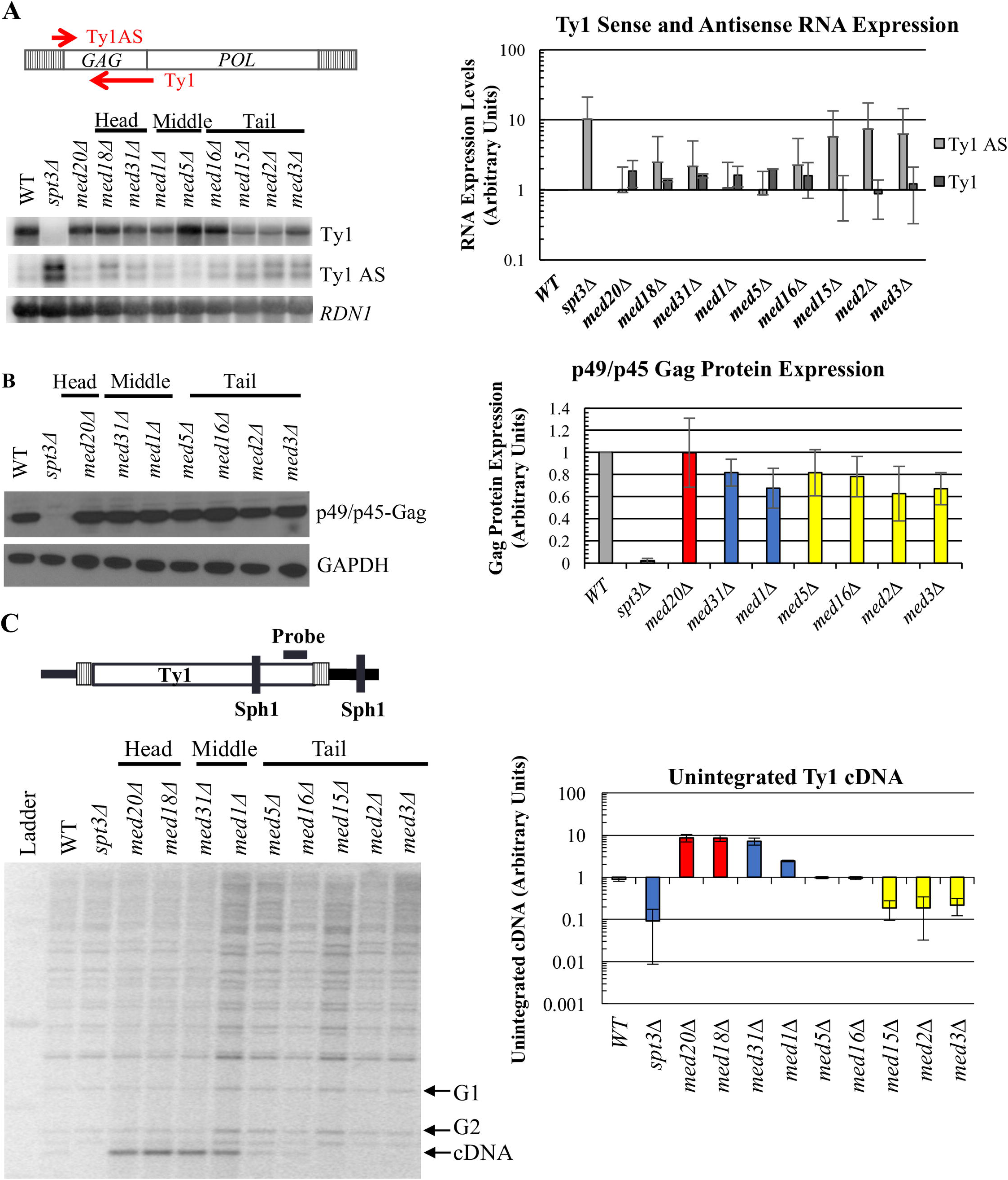
Mediator subunits influence Ty1 cDNA levels without altering levels of Ty1 RNA or Gag protein. (A) Quantitative northern blot analysis of sense-strand (Ty1) and antisense-strand (Ty1 AS) RNA. Total RNA was fractioned by gel electrophoresis, blotted, and the membranes probed for Ty1 AS RNA, followed by stripping and probing for Ty1 RNA using strand-specific riboprobes as schematized at the top, and for 18S rRNA as a loading control. The graph on the right shows the quantitation of Ty1 RNA and Ty1 AS RNA levels from two biological replicates, each relative to the 18S subunit rRNA level (note log scale), normalized to WT levels. (B) Western blot of total cell lysates probed for Gag using a polyclonal antibody against p18 [24]. Data for *med5Δ* and *med16Δ* is derived from two biological replicates; all other values represent data from three biological replicates, and values are normalized to WT. (C) Quantitative Southern blot analysis to determine the level of unintegrated Ty1 cDNA (cDNA) relative to the amount of DNA in bands representing two genomic Ty1 elements (G1 and G2). Image quantification (note log scale) is representative of three biological replicates, only one of which is shown. All experiments were performed using congenic WT, *spt3A* and Mediator subunit deletion strains harboring *Ty1his3AI-3*114. Bars in (B) and (C) are color coded as in Figure 2, and all error bars represent s.d. Note log scale.

A series of non-coding antisense transcripts are also expressed from Ty1 elements via an internal promoter (Fig 1B & C). These transcripts initiate from distinct positions within the first 700 bp of the Ty1 element, and their expression is enhanced in certain hypomobile strains, most notably in an *spt3Δ* mutant [27, 31]. Levels of Ty1 antisense transcripts (Ty1AS RNA) were measured to determine whether Mediator was altering expression of these transcripts and thereby influencing Ty1 mobility (Fig 3A). A modest increase of approximately 2-fold in Ty1AS RNA was observed in the Mediator tail subunit deletion strains relative to the wild-type strain (Fig 3A). An increase of a similar magnitude was observed in the hypermobile *med31Δ* strain, while a 10-fold increase was measured in the hypomobile *spt3A* strain. Based on the lack of correlation with altered retromobility, Ty1AS RNA expression is not likely to be a major contributing factor to tail-mediated repression of Ty1 retromobility.

To independently confirm that the substantial changes in retromobility in Mediator subunit deletion strains were not a result of minor changes in Ty1 RNA levels, or in alterations to Ty1 polyadenylation that would result in translational defects, the levels of Gag, the major product of Ty1 RNA translation, were measured using western blot analysis (Fig 3B and S1 Fig). As with Ty1 RNA and Ty1AS RNA, there were only moderate (<2-fold) changes in the levels of Gag protein in Mediator subunit deletion strains relative to the wild-type strain. Together, these data support the argument that Mediator regulates a post-transcriptional step in Ty1 retromobility, and does so without altering the steady-state level of Gag (Fig 3A & B).

Assessing the level of unintegrated cDNA provides an indication of the efficiency with which Ty1 proteins and RNA have assembled into functional VLPs and carried out reverse transcription. This assay involves the electrophoretic separation of SphI-digested genomic DNA, which is subsequently probed for Ty1 sequences at the 3’ end of *POL* [15, 37], allowing for the visualization of bands representing the junction between the 3’ end of each Ty1 element and flanking genomic DNA. Differences in the size of the bands are due to the different location of *SphI* sites in DNA flanking Ty1 at different locations. In this assay, the smallest band represents unintegrated Ty1 cDNA because of the absence of flanking genomic DNA. We performed this assay to compare the ratio of unintegrated cDNA to genomic Ty1 DNA in wild-type yeast to that in Mediator mutants. We observed increased cDNA levels for *Med18Δ*, *Med20Δ*, *med31Δ*, and *Med1Δ* mutants, while loss of tail module triad subunits Med2, Med3, or Med15 reduced cDNA to nearly undetectable levels (Fig 3C). Thus, both hyper- and hypomobile mutants exhibited cDNA levels that correlated well with changes in Ty1 *his3AI* mobility (Fig 2C). The magnitude of changes in Ty1 cDNA levels are not as great as those of Ty1 retromobility because the cDNA assay measures steady-state levels, whereas the retromobility assay measures accumulated events. Together, these data indicate that Mediator core subunits regulate a post-transcriptional step in Ty1 mobility, such that deletion of Mediator complex genes alters the accumulation of Ty1 cDNA without substantially altering overall Ty1 transcript or Gag protein levels.

### The Mediator Tail Module Regulates Ty1 Retromobility in cis via an Interaction with the LTR Promoter

Mediator mutants might affect Ty1 activity indirectly by altering the expression of a host factor(s) that controls a post-transcriptional step in retrotransposition. If this were the case, similar effects on retromobility would be observed whether Ty1*his3AI* RNA was expressed from the LTR promoter or a heterologous promoter. Alternatively, Mediator mutants could affect Ty1 activity in *cis* by acting at the genomic Ty1 promoter or an internal Ty1 promoter. In this case, the effects might be suppressed by expression of Ty1*his3AI* RNA from a heterologous promoter.

To differentiate between these alternatives, the Ty1 promoter in the U3 region of the LTR was swapped for the transcriptionally robust *TEF1* promoter sequence in a CEN-plasmid-based Ty1*his3AI* element (Fig 4A). The *TEF1* promoter was chosen based on findings that deletions of non-essential Mediator subunits do not alter *TEF1* expression substantially [59, 60]. The effects of Mediator subunit deletions on retromobility of the *TEF1* promoter-driven Ty1*his3AI* element on a CEN-plasmid versus the LTR promoter-driven Ty1*his3AI* element on a CEN-plasmid were then compared (Fig 4B).

**Fig 4:**
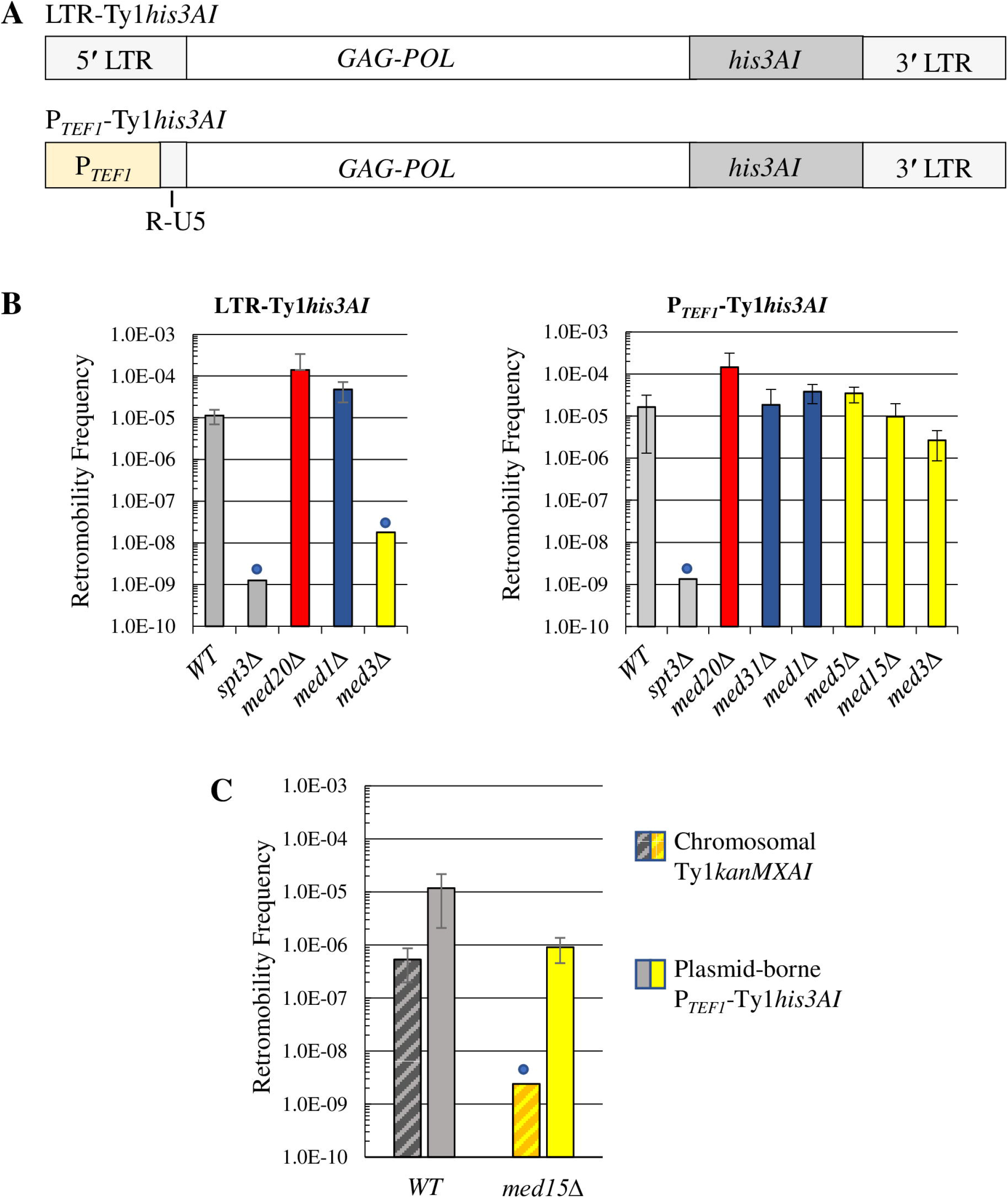
The Mediator tail acts on Ty1 mobility in an LTR promoter-dependent manner. (A) Schematic of the P_*TEF1*_-Ty1*his3AI* element relative to a Ty1*his3AI* element with the standard LTR promoter. P_*TEF1*_-Ty1*his3AI* has a *TEF1* promoter in place of Ty1 promoter elements in the U3 region of the 5’ LTR (See Fig 1C), while retaining the Ty1 TSS and R-U5 region of the 5’ LTR. (B) Retrotransposition frequency, shown on a log scale, for a plasmid-based Ty1*his3AI* (left) or P *TEFı-* Ty1*his3AI* (right) element in the WT and *spt3A* negative control strains (grey bars), Mediator head subunit gene deletion strains (red bars), middle subunit gene deletion strains (blue bars) and tail subunit gene deletion strains (yellow bars). (C) Retromobility frequencies, shown on a log scale, of a chromosomal Ty1*kanMXAI* element and a plasmid-borne P_*TEF1*_-Ty1*his3AI* element contained in the same wild type or *Med15Δ* strain. All error bars represent s.d. Blue circles in (B) and (C) denote values that represent upper limit mobility estimates for strains in which most or all cultures had no His+ prototrophs (LTR-Ty1 *his3AI*or *P_TE_Fi-Ty1*his3AI*)* or G418^R^ colonies (Ty1*kanMXAI*) per total number of Ura+ or Leu+ cells analyzed, respectively.

Deletion of the tail module triad gene *MED3* reduced retromobility of the plasmid-borne Ty1*his3AI* element more than 600-fold (Fig. 4B), which is consistent with the effect of this deletion on a chromosomal Ty1*his3AI* element (Fig. 2C). In contrast, retromobility of the *P_*TEF1*_*-driven Ty1*his3AI* element was reduced 6-fold or less by deletion of *MED3* or *MED15* (Fig 4B). Together, these results indicate that the Mediator tail triad module controls Ty1*his3AI* retromobility in an LTR-dependent fashion. Middle or head module gene deletions increased retromobility of the plasmid-borne Ty1*his3AI* element to a lesser extent than that of the chromosomal Ty1*his3AI* element (≤12-fold versus >100-fold; compare Fig. 4B and Fig 2), possibly because retromobility of Ty1*his3AI* on the CEN-plasmid is already substantially higher than that of the chromosomal element. Correspondingly, deletion of the head subunit gene *MED20* or middle subunit gene *MED31* or *MED1* increased retromobility of the plasmid-driven P_*TEF1*_-Ty1*his3AI* and Ty1*his3AI* elements to a similar extent (Fig. 4B). Consequently, we could not determine from these data whether Mediator head and middle subunits regulate Ty1*his3AI* retromobility in *cis* or in *trans*.

Regulation of retromobility by the Mediator tail module triad when Ty1*his3AI* RNA is driven from the LTR versus the TEF1 promoter suggests that Mediator acts in *cis* on Ty1*his3AI* To further test this idea, the CEN-plasmid bearing P_*TEF1*_-Ty1*his3AI* was introduced into congenic wild-type and *Med15Δ* strains that harbor a chromosomal Ty1*kanMXAI* element. The effect of the *Med15Δ* deletion on retromobility of the chromosomal Ty1*kanMXAI* element, which is measured by determining the frequency of G418^R^ prototroph formation, was compared to its effect on retromobility of the plasmid-borne P_*TEF1*_-Ty1*his3AI* element (Fig. 4C). Deleting *MED15* had a modest effect on P_*TEF1*_-Ty1*his3AI* retromobility, but it reduced Ty1*kanMXAI* retromobility >200-fold to a level below detection. The different effects of deleting *MED15* on these elements could reflect greater suppression of the hybrid *TEF1* promoter than the LTR promoter by the Mediator tail module triad; indeed, P_*TEF1*_-driven Ty1*his3AI* RNA is selectively and significantly increased in a *Med15Δ* mutant (S2 Fig). Taken together, these data suggest that Mediator, via its tail module triad, functions directly at *Ty1*his3AI*to* enhance retromobility by a mechanism that depends at least partially on the U3 region of the Ty1 LTR.

### The Tail Module Triad Regulates Ty1 via Modulation of Ty1i Expression

The observation that the Mediator tail module triad is a potent regulator of retromobility that functions in *cis* at Ty1 without significantly influencing Ty1 RNA levels led us to consider the possibility that Mediator regulates the balance between the expression of the internal Ty1i transcript, which encodes a dominant negative inhibitor of retromobility, and expression of the genomic Ty1 transcript. The post-transcriptional effects on Ty1 retromobility caused by increasing Ty1i expression [24] are consistent with those observed in Mediator tail triad deletion mutants. We therefore sought to determine whether expression of Ty1i RNA is altered by deletion of Mediator subunits.

Ty1i RNA is not easily detected in northern blots of total cellular mRNA (Fig 3A), as it is obscured by the highly abundant Ty1 transcript [24]. Only about 15% of Ty1 RNA is polyadenylated [35]; therefore polyA+ RNA of the wild-type strain and Mediator deletion strains was subjected to northern analysis to achieve better separation of the Ty1 and Ty1i transcripts, as previously reported [24]. This analysis revealed an increase in Ty1i RNA in all three mutants with a tail module triad gene deletion, consistent with reduced retromobility in these mutants, while *Med5Δ* yeast, which showed no change in retromobility, had Ty1i RNA levels similar to those observed in wild-type yeast (Fig 5A). In contrast, the level of genomic Ty1 RNA was altered only minimally by deletion of a tail module triad gene or *MED5.* Neither the level of Ty1i RNA nor the level of Ty1 RNA in head and middle subunit deletion strains was substantially altered relative to those in the wild-type strain (Fig 5A). Given the low level of Ty1i RNA in the wild-type strain, we were unable to determine whether deletion of a head or middle subunit results in a decrease in Ty1i RNA that could account for the hypermobility phenotypes of these mutants.

**Fig 5:**
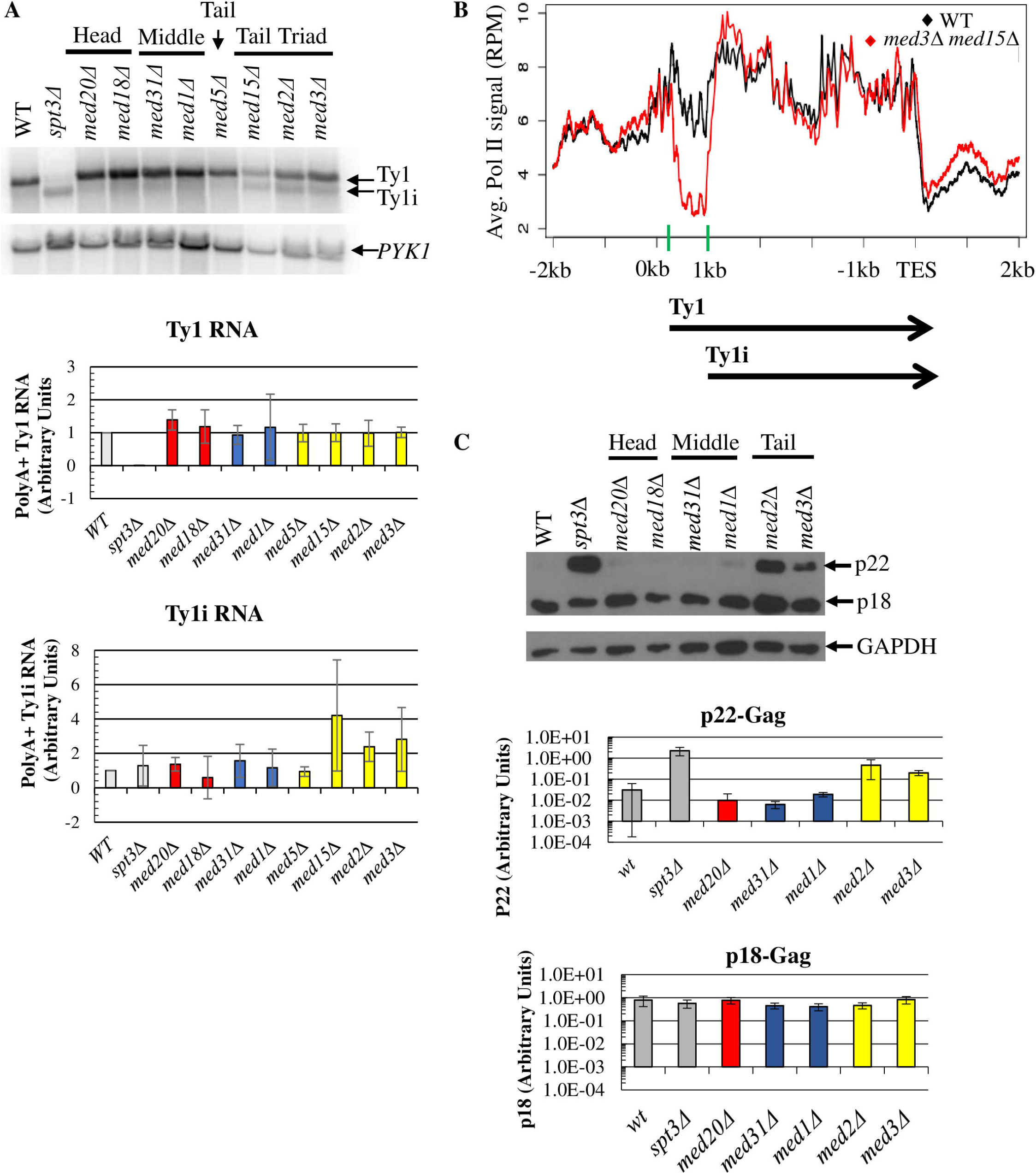
Deletion of Mediator tail module triad subunits increases levels of polyA+ Tyli RNA and p22-Gag. (A) Northern blot probed for Ty1, Ty1i and *PYK1* RNA, the latter as a loading control that has been shown not to be altered in *Med2Δ*, *Med3Δ*, or *Med15Δ* yeast [57, 60]. Quantifications of Ty1 and Ty1i RNA relative to *PYK1* RNA are averages of three biological replicates, except for *Med18Δ*, which is an average of two biological replicates, and is normalized to WT levels. (B) Pol II occupancy averaged over all 31 genomic Ty1 elements in wild type and *Med3Δ Med15Δ* yeast using ChIP-seq data from [61]. Ty1 elements begin at 0 kb on the x-axis, and the Ty1 TSS at +238 and Ty1i TSS at +1000 are marked by green bars on the x-axis. (C) Western blot of total cell lysate measuring levels of p22- and p18-Gag relative to the loading control, GAPDH.. Quantitation, shown on a log scale, is the average ratio of p22-Gag or p18-Gag to GAPDH signal from three biological replicates of each strain. In panels (A) and (C), quantitation was performed on RNA or protein samples from the WT strain and congenic *spt3A* strain as a negative control (grey bars), Mediator head subunit gene deletion strains (red bars), middle subunit gene deletion strains (blue bars) and tail subunit gene deletion strains (yellow bars). All error bars represent s.d.

Previous work from the Morse lab compared genome-wide occupancy of Pol II in wild type and *Med3Δ Med15Δ* yeast, although not in single tail module triad gene deletion mutants [61]. The effects of the *Med3Δ Med15Δ* mutation on growth and genome-wide transcription are similar to those of the single Mediator tail module triad subunit deletions [57], and so we used this data to compare Pol II occupancy at all Ty1 elements in wild type and *Med3Δ Med15Δ* yeast (Fig 5B). The results show that Pol II occupancy in *Med3Δ Med15Δ* yeast is reduced exclusively in the first 1 kb of the Ty1 element. Pol II occupancy in the *Med3Δ med15Δ* double mutant and wild-type strain became equivalent near the Ty1i transcription start site (TSS) and remained so until the transcription end site (TES) of both Ty1 sense-strand transcripts (Fig 5B). (Close inspection of Fig. 5B shows that Pol II ChIP signal at the Ty1i TSS region is actually steeply increasing in *med3 med15* yeast, reaching levels equivalent to those seen in wild type yeast at about +200 relative to this TSS. This is consistent with occupancy becoming equivalent close to the TSS, because the fragments analyzed in ChIP experiments vary from 200 to about 500 bp in length, so that decreased occupancy at a particular location results in decreased ChIP signal on either side of that location [62]). The decreased occupancy by Pol II over the first ~700 bp of Ty1 in yeast lacking tail module triad subunits indicates that the Mediator tail module plays a critical role in establishing the relative occupancy of Pol II at the Ty1 and Ty1i TSS, consistent with the altered ratio of Ty1i RNA to Ty1 RNA levels observed in tail module triad mutants (Fig 5A).

The elevated Ty1i RNA levels observed in Mediator tail module triad mutants would be predicted to give rise to increased levels of the retromobility inhibitor, p22-Gag and its cleavage product, p18-Gag. To test this prediction, we performed western blotting using an anti-p18 polyclonal antibody [24]. Elevated levels of p22-Gag were observed in *Med2Δ* and *Med3Δ* yeast, in accord with the increased Ty1i levels in these mutants, whereas virtually no p22-Gag was detected in the wild-type strain or Mediator head and middle module mutants (Fig 5C). Unexpectedly and in contrast to previous observations in *Saccharomyces paradoxus* [24], levels of p18-Gag were similar in all strains and were not correlated with p22-Gag or Ty1 retromobility levels in the wild-type strain or any Mediator subunit deletion strain. Nonetheless, these data demonstrate that diminished Ty1 retromobility is accompanied by elevated Ty1i RNA and p22-Gag levels in Mediator tail module mutants. Taken together with the increased occupancy of Pol II at Ty1i relative to Ty1 in *Med3Δ Med15Δ* yeast (Fig 5B) and previous findings that p22-Gag blocks post-transcriptional steps in Ty1 mobility, these results suggest that loss of Mediator tail module triad subunits blocks Ty1 retrotransposition at a post-transcriptional step by causing increased Pol II recruitment to the Ty1i promoter relative to the Ty1 promoter.

### Mediator Tail and Head Module Subunits Direct Mediator Association with Ty1 and Ty1i Proximal Promoters, Respectively

Results presented so far indicate that the opposing effects of deletion of subunits from the Mediator head module (*med18Δ* and *med20Δ*) and tail module triad (*med2Δ, med3Δ*, and *med15Δ*) on Ty1 mobility occur via mechanisms that operate at a similar stage in the Ty1 life cycle (Fig 3). Decreased mobility in mutants lacking subunits from the tail module triad is accompanied by increased expression of Ty1i, and similarly decreased occupancy of Pol II in the first 1kb of Ty1, suggesting a causal mechanism (Fig 5). However, Ty1i RNA and p22-Gag levels are very low in wild type yeast, and we have not been able to detect any decrease of this already low level in *med18Δ* or *med20Δ* mutants.

To investigate the mechanism by which tail module triad and head module deletions exert opposing effects on Ty1 mobility, we used ChIP-seq to examine Mediator association with the proximal promoter regions of Ty1 and Ty1i in wild type yeast and Mediator mutants. For this purpose, we used *kin28* Anchor Away (*kin28-AA*) strains harboring appropriate deletions of Mediator subunits. The *kin28-AA* conditional mutation allows eviction of Kin28 from the nucleus to the cytoplasm by addition of rapamycin [63, 64]. Eviction of Kin28 prevents phosphorylation of Ser5 of the Pol II C-terminal domain (CTD). This impedes release of engaged Pol II from the proximal promoter of active genes, thus stabilizing association of Mediator, allowing robust ChIP signals to be observed at gene promoters [64, 65].

ChIP-seq against myc-tagged Med17 from the head module or Med15 from the tail module reveals peaks of Mediator association at both Ty1 and Ty1i TSS in wild type yeast (Fig 6 and S3 Fig). In a *med2Δ med3Δ med15Δ* mutant, Med17 association with the Ty1 TSS is decreased relative to its association with the Ty1i TSS (Fig 6), consistent with increased Ty1i RNA levels and increased occupancy by Pol II at Ty1i compared to Ty1 (Fig 5B and S4 Fig) in tail module triad deletion mutants. In contrast, occupancy by Med17 (from the head module) and Med15 (tail) at Ty1i is virtually eliminated in *Med18Δ* (S3 Fig) and *Med20Δ* (Fig 6) mutants, while Ty1 occupancy remains robust. Western blotting showed that Med17-myc expression was about 1.6-fold greater in *Med18Δ* and *Med20Δ* mutants and about 2.3-fold greater in *med2Δ med3Δ med15Δ* yeast than in wild type yeast (S5 Fig). We were unable to reliably detect Med15-myc in western blots, possibly due to poor transfer of this large protein. These results suggest that the relative occupancy levels by Mediator, and presumably by the PIC, at Ty1 and Ty1i proximal promoters are dictated by Mediator subunits, and play a critical role in governing Ty1 mobility.

**Fig 6:**
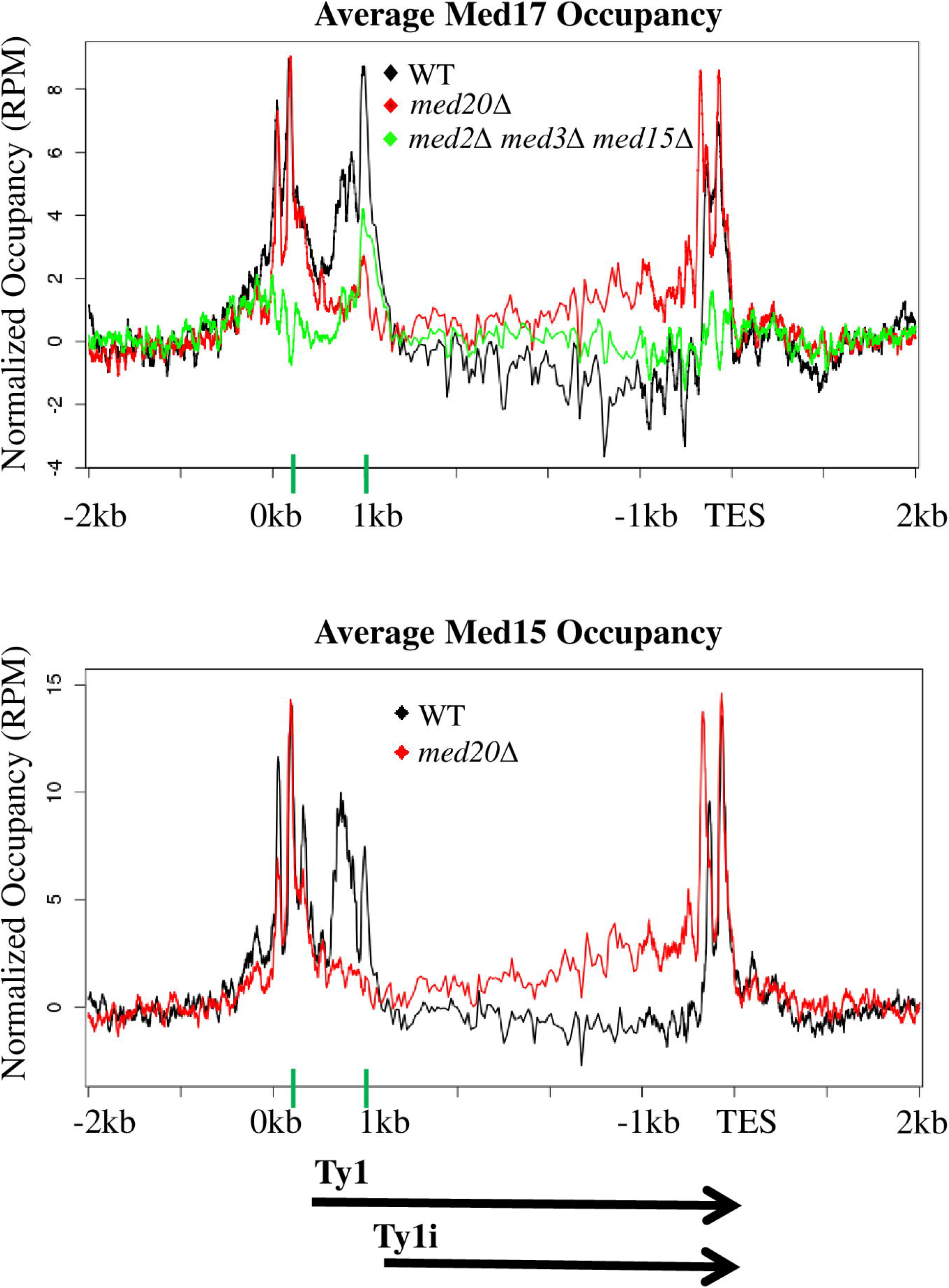
Mediator occupancy at the Ty1 and Ty1i proximal promoters. Occupancy of Med17-myc and Med15-myc from the Mediator head and tail modules, respectively, was determined by ChIP-seq and summed over all Ty1 elements in *kin28-AA* yeast that were otherwise wild type (WT), *Med20Δ*, or carried the triple deletion *Med2Δ Med3Δ Med15Δ*. Ty1 elements begin at 0 kb on the x-axis, and the Ty1 TSS at +238 and Ty1i TSS at +1000 are marked by green bars on the x-axis. ChIP-seq signals were normalized to an untagged *kin28-AA* control (S3 Fig). Reads deriving from the Ty1 TSS, and therefore in the LTR, are unavoidably also assigned to the 3’ LTR, leading to the observed signal in that region.

### Ty1i RNA Is Repressed by the Mediator Tail Module Triad Independently of Full Length Ty1 Transcription

Our results indicate that the presence of the Mediator tail module triad prevents expression of Ty1i RNA. We considered two mechanisms by which this might occur. First, tail-module dependent recruitment of Mediator to the Ty1 promoter could lead to enhanced Ty1 transcription, and this could repress the downstream Ty1i promoter by read-through effects. In this mechanism, the polymerase moving from the Ty1 TSS could disrupt transcription factor or PIC binding to the Ty1i promoter [66-68]. In the second mechanism, the Mediator tail could act as a direct repressor of the Ty1i promoter.

To distinguish between these two mechanistic possibilities, we asked whether increased Ty1i expression was still observed in tail module triad deletion mutants under conditions where transcription from the Ty1 promoter was strongly repressed. To this end, we employed a plasmid-based Ty1 element under control of the *GAL1* promoter, which is strongly repressed in glucose medium (Fig 7) [69]. The 2μM-plasmid-borne *GAL1*-Ty1 element (pGTy1) that we used also has an internal deletion within the *POL* ORF, which facilitates resolution of Ty1 ΔPOL RNA from genomic Ty1 mRNA, thus permitting a direct comparison between endogenous Ty1 RNA and plasmid-derived Ty1 ΔPOL RNA by northern blot or single strand cDNA synthesis followed by PCR analysis.

**Fig 7:**
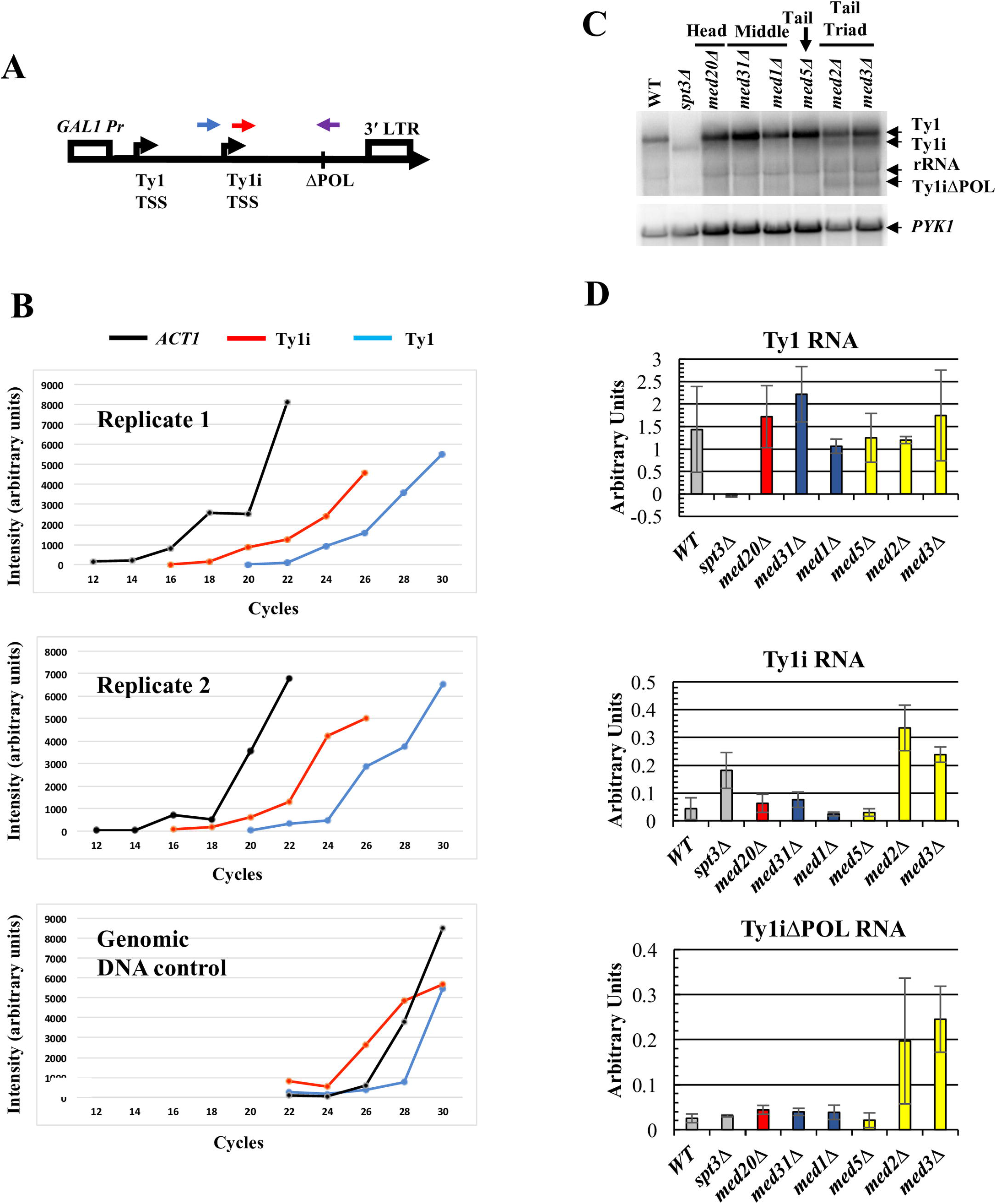
Ty1i situated downstream of a repressed Ty1 element is repressed by Mediator tail module triad subunits. (A) Schematic of the GAL1:Ty1ΔPOL cassette in pGTy1ΔPOL showing forward primer locations for detection of Ty1 RNA (blue) versus Ty1i RNA (red). Note that because Ty1i is contained within Ty1, the Ty1i primer reports both Ty1i and Ty1 transcripts. Both amplifications utilized the same reverse primer (purple), that crosses the deletion junction and contains sequences unique to the pGTy1ΔPOL element. A reverse primer specific for *ACT1* mRNA was also used to synthesize cDNA used as a template for the PCR amplification. No PCR product was detected using the POL reverse primer when RNA from yeast lacking pGTy1ΔPOL was used as template. (B) Quantitation of the products of Reverse Transciption-PCR reactions using polyA+ RNA isolated from wild-type yeast bearing plasmid pGTy1ΔPOL, and grown in glucose-containing broth, and from a genomic DNA control. Aliquots were taken from reactions at the indicated number of cycles and analyzed by agarose gel electrophoresis (S6 Fig). Levels of RT-PCR amplification products using Ty1, Ty1i, and *ACT1* primers are indicated. (C) Northern blot probed with a single-stranded riboprobe that hybridizes to Ty1, Ty1i, Ty1ΔPOL and Ty1iΔPOL RNA. Note that the full-length Ty1ΔPOL transcript cannot be distinguished from the rRNA band. Image is representative of three biological replicates. (D) Quantification for Ty1, Ty1i, and Ty1iΔPOL RNA from three biological replicate northern analyses. Graph bar colors correspond to WT strain and the congenic *spt3A* strain as a negative control (grey bars), Mediator head subunit gene deletion strain (red bars), middle subunit gene deletion strains (blue bars) and tail subunit gene deletion strains (yellow bars). Error bars represent s.d.

We confirmed that Ty1ΔPOL and Ty1iΔPOL RNAs are expressed at very low levels in cells grown in glucose by using a PCR assay that distinguishes these transcripts from endogenous Ty1 RNA. To this end, a reverse primer spanning the ΔPOL deletion was used with either of two forward primers located 250bp upstream or 250bp downstream of the Ty1i TSS (Fig 7, top). The upstream (blue in Fig. 7) primer amplifies only Ty1ΔPOL cDNA, while the Ty1i primer (red in Fig. 7) amplifies both Ty1 and Ty1i products. Use of these primers precluded use of real time PCR; instead, aliquots from the reactions were removed at two cycle intervals for analysis by gel electrophoresis (S6 Fig). Results from two replicate experiments using this strategy are depicted in Figure 7B. Taking into account the differential amplification observed using a genomic DNA control (Figure 7B, bottom panel, and S6 Fig), Ty1ΔPOL transcript levels appear lower than or comparable to levels of Ty1ΔPOL plus Ty1iΔPOL transcripts. More importantly, we estimate that these transcripts are present at less than 1% of the abundance of *ACT1* mRNA (see Methods). We conclude that pGTy1ΔPOL is transcribed at very low levels in cells grown in glucose medium, as expected.

To examine the effect of Mediator subunit deletions on Ty1i RNA expression from pGTy1ΔPOL, levels of Ty1 and Ty1i RNA from endogenous Ty1 elements and Ty1ΔPOL and Ty1iΔPOL RNA from pGTy1ΔPOL were measured by northern blotting (Fig 7C-D). Deletion of Mediator subunits had little effect on levels of endogenous genomic Ty1 RNA, as expected. Ty1i and Ty1iΔPOL transcripts were barely detectable in the wild-type strain and head and middle subunit deletion strains; however, Ty1i and Ty1iΔPOL RNAs were markedly increased in strains lacking the tail module subunits, *MED2* or *MED3* (Fig 7C-D). Therefore, disruption of the Mediator tail increases Ty1iΔPOL RNA expression, even when expression from the upstream Ty1 promoter is repressed. This indicates that the tail module triad does not suppress expression of Ty1i RNA via read-through inhibition from the upstream Ty1 promoter. Taken together with the altered occupancy of Pol II and Mediator over Ty1 seen in yeast lacking tail module triad subunits (Fig 5B and Fig 6), these results suggest that the Mediator tail module triad acts to direct PIC formation preferentially to the Ty1 TSS over the Ty1i TSS.

## Discussion

Previous studies provided evidence that loss of various non-essential Mediator subunits affected Ty1 mobility [34-39, 53], but they did not systematically characterize the role of Mediator in this process, nor did they provide an explanation for how a transcriptional activator could exert its effects post-transcriptionally. By examining the effects of deletion of each non-essential Mediator subunit on Ty1 retromobility, we show that Mediator functions as both an activator and repressor of Ty1 retromobility (Fig 2B). Effects of Mediator subunit deletions on retromobility correlated well with alterations in Ty1 cDNA levels, but not with changes in Ty1 RNA levels or Gag protein levels, as shown previously for some subunit deletions [35-38]. These findings led us to examine the effect of Mediator subunit deletions on expression of Ty1i, which encodes a dominant inhibitor of Ty1 cDNA synthesis [24]. We found that Ty1i expression was increased in mutants lacking tail module triad subunits, in which retrotransposition is reduced to undetectable levels. Relative occupancy of Pol II at Ty1 and Ty1i in *med3Δ med15Δ* yeast was altered in favor of Ty1i, and Mediator occupancy in yeast lacking tail module triad subunits was also altered to favor the Ty1i promoter. Taken together, these findings indicate that loss of tail module triad subunits alters the balance of utilization of Ty1 and Ty1i promoters, resulting in increased Ty1i/Ty1 RNA and concomitant inhibition of retromobility.

Deletion of head and middle module subunits had opposite effects to those seen with tail module triad subunit deletions, namely, increased retrotransposition and Ty1 cDNA levels. It thus seemed likely that these mutations also exerted their effects by altering the balance of expression between Ty1 and Ty1i. The low levels of Ty1i present in wild type cells made determination of any reduction in these levels challenging; furthermore, only a very small fraction of a given population of yeast cells undergo retrotransposition, and currently there is no way to assess the molecular properties of that specific subpopulation. Nonetheless, we were able to observe altered Mediator occupancy in favor of Ty1 relative to Ty1i in *med18Δ* and *med20Δ* mutants, supporting the idea of a common mechanism.

These findings are summarized in the model depicted in Fig 8. The Ty1 LTR contains a TATA box within its U3 region (Fig 1), and Ty1 expression is Spt3-dependent (Fig 3 & 5) [25], indicating that Ty1 belongs to the SAGA-dominated class of genes. This class is enriched in highly regulated genes such as stress-response genes, and is characterized by a promoter structure that is distinct from that of the largely constitutively-active, TATA-less, TFIID-dominated genes [20, 70]. In contrast to Ty1, Ty1i is transcribed in *spt3Δ* yeast and is therefore not dependent on SAGA; furthermore, the region upstream of the Ty1i TSS lacks any consensus TATA element, indicating that Ty1i belongs to the class of TFIID-dominated genes. Genes whose activation depends on the Mediator tail module triad are relatively enriched for TATA-containing, SAGA-dominated family members [57]; in accordance with this observation, we propose that mutants lacking tail module triad subunits have reduced efficiency of PIC formation at the Ty1 promoter, and that this permits increased utilization of the TATA-less Ty1i promoter. Conversely, we propose that the TATA-less Ty1i promoter depends on head and middle module subunits for Mediator association and PIC formation, consistent with our Mediator ChIP-seq results. Finally, we suggest that replacing the TATA-containing Ty1 promoter with the strong, TFIID-dominated *TEF1* promoter allows Ty1 transcription to successfully compete against the Ty1i promoter even when tail module triad subunits are deleted, so that no increase in *Ty1ihis3AI* RNA is observed for this reporter (S2 Fig).

**Fig 8:**
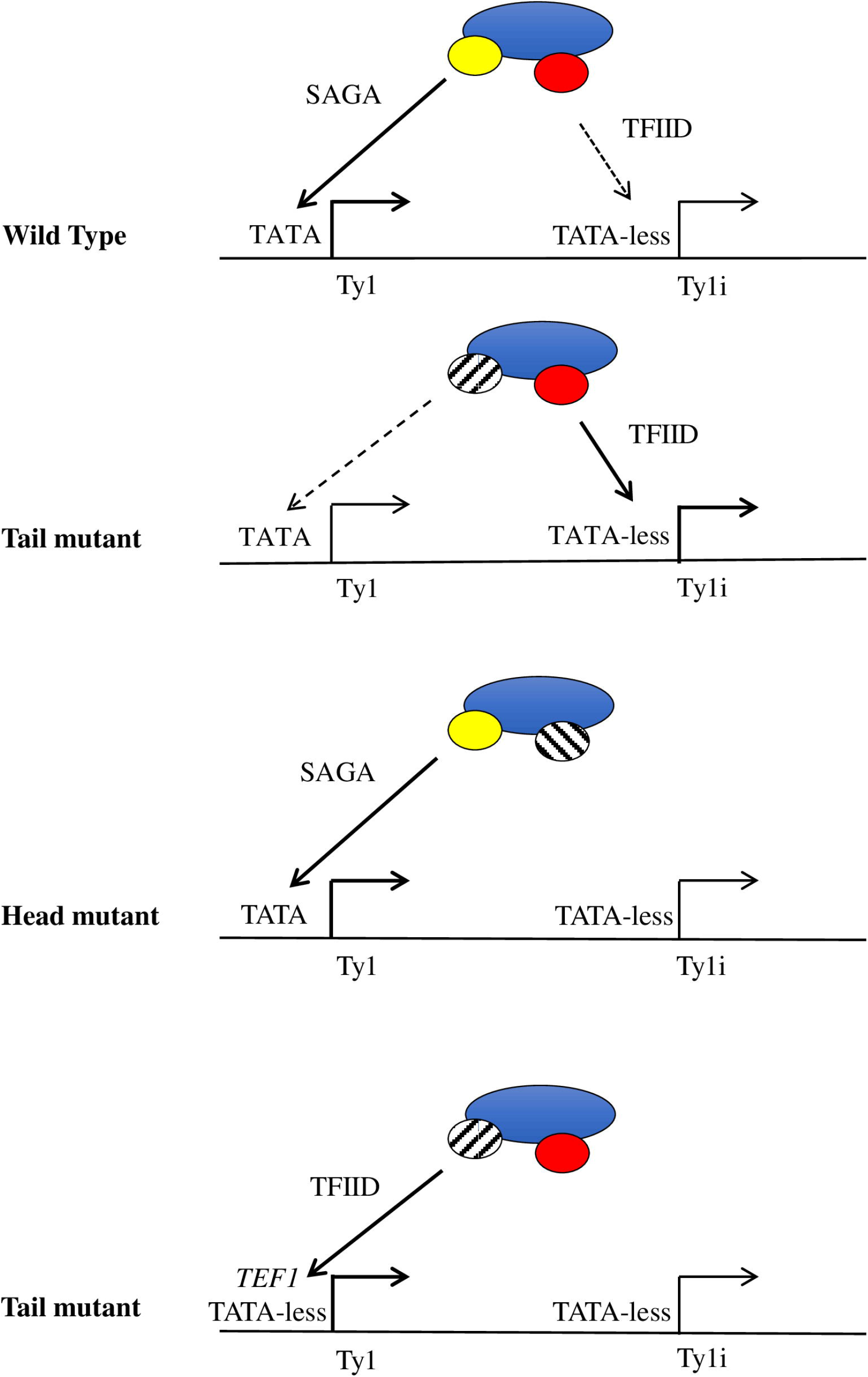
Model for altered balance of utilization of Ty1 and Ty1i promoters in Mediator deletion mutants. In wild type yeast, the Mediator complex acts at both the Ty1 and Ty1i proximal promoters, stimulating robust transcription of Ty1 and a small amount of Ty1i expression that is sufficient to restrict mobility. Deletion of subunits from the tail module triad increases Mediator activity at the Ty1i promoter, thus increasing Ty1i production and reducing retromobility. Conversely, when a subunit from the head module is deleted, the association between Mediator and the Ty1i promoter is perturbed, permitting increases in retromobility. Finally, when the SAGA-dependent Ty1 promoter is swapped for the strong, TFIID-dependent *TEF1* promoter, the complex preferentially associates with the Ty1 promoter, irrespective of the influence of the tail module.

Our model and our ChIP-seq results indicate substantial Mediator occupancy at both the Ty1 and the Ty1i promoter in wild-type yeast, despite the apparent differences in transcriptional output from these two promoters. There are several possible explanations for this disparity. First, Mediator occupancy was measured under conditions of Kin28 depletion, potentially obscuring any differences caused by different efficiencies in facilitating PIC formation and productive transcription at the two promoters. Second, Ty1i RNA may be substantially less stable than Ty1 RNA. Indeed, Ty1 RNA is known to be unusually stable [71, 72]. Third, transcription from Ty1i may be inefficient in spite of the apparent presence of Mediator at the Ty1i proximal promoter. This may be altered in Mediator head and middle module mutants, or it may be that a minor change in Ty1i RNA abundance leads to a significant change in the small population of cells that undergo transposition. It is also possible that the head and middle modules additionally influence other steps in the retrotransposition life cycle. For example, loss of head and middle module subunits may alter Ty1 RNA localization in such a way as to prevent retrotransposition. This possibility is supported by observed increases in ribonucleoprotein foci known as retrosomes in *med20Δ*, *med18Δ*, *med9Δ*, and *med1Δ* mutants [35].

The molecular mechanism underlying the contrasting dependence of Ty1 and Ty1i promoters on Mediator subunits remains to be determined, but is likely to involve Ty1 promoter elements that are present upstream of the Ty1i proximal promoter. Ty1 possesses an unusual promoter architecture that includes transcription factor binding sites both upstream of the Ty1 TSS and sites within the ORF, but upstream of the Ty1i TSS (Fig 1C) [18]. The latter region includes sites for Ste12, Tec1, Tye7, Mcm1, Rap1, and Tea1; intriguingly, Med15 has been shown to negatively regulate activation of a reporter gene by this intra-ORF region via the Mcm1 site [73]. Conceivably, interaction of Mediator with this region may be sufficient to inhibit PIC formation at the Ty1i proximal promoter, in a manner dependent on the tail module triad, even when the Ty1 proximal promoter is inactive, as is the case for pGTy1ΔPOL in yeast grown in glucose medium.

Retroelements are nearly ubiquitious in extant genomes, and their expression is governed by disparate mechanisms, including expression of truncated ORFs and altered TSS utilization [74, 75]. We are not aware of any precedent for Mediator regulating the balance between two promoters in the way we have reported here; future studies will be required to determine whether similar mechanisms apply in other cases in yeast or other organisms. Regarding physiological relevance, retrotransposition frequency is responsive to environmental stress, and Mediator is subject to stress-dependent phosphorylation that affects gene expression [76]. This suggests a possible mechanism for regulating Ty1 retromobility during stress that deserves exploration.

An unanticipated finding of this study was that retromobility of an LTR-driven Ty1*his3AI* element on a low copy CEN-plasmid was over 30 times higher than that of the active chromosomal element, *Ty1*his3AI*-3*114 (Compare Fig 2B to Fig 4B). Because the amount of Ty1*his3AI*RNA relative to total Ty1 RNA is a direct determinant of the frequency of Ty1*his3AI* retromobility, these data may indicate that the LTR promoter is more active on the CEN-plasmid, possibly due to reduced nucleosome occupancy at the LTR [77, 78]. Potential differences in Ty1 or Ty1i RNA expression between chromosomal and CEN-plasmid elements warrants further investigation, as these might explain why different screens for Ty1 regulators have yielded largely non-overlapping gene sets [18].

A frequent finding among studies of extrinsic and intrinsic regulators of Ty1 is that the retromobility level is altered without a change in the level of Gag protein [18]. For example, treatment of cells with the DNA damaging agents 4-Nitroquinoline 1-oxide and γ-rays, severe adenine starvation, reduced growth temperature and telomere erosion in the absence of telomerase induce Ty1 retromobility, and in some cases, increase Ty1 RNA without altering steady-state Gag levels [18, 79-81].

In addition, treatment of cells with γ-factor, the absence of 5’ to 3’ mRNA degradation proteins, and increased Ty1 copy number all restrict Ty1 mobility with minor if any effects on Gag protein levels [15, 24, 82, 83]. Increased Ty1 copy number and the absence of 5’ to 3’ mRNA decay factors leads to increased Ty1i RNA expression [24, 31, 33], but the role of Ty1i RNA in these other phenomena has not yet been determined. Perhaps the Mediator complex and Ty1i/Ty1 promoter balance play a role in these regulatory processes. *Saccharomyces cerevisiae* lacks RNA interference [84], which invites speculation that Mediator has assumed the function of central coordinator of Ty1 retrotransposon activity by integrating diverse extrinsic and intrinsic signals and modulating the balance between Ty1i and Ty1 expression.

## Materials and Methods

### Yeast Strains and Plasmids

Strains used in this study are derivatives of BY4741. Genotypes of each strain are provided in S1 Table. Strains containing a chromosomal *his3AI*-[Δ*1*]-marked Ty1 element (Ty1*his3AI*-3114) were described previously [19, 85]. Recombination of the *his3AI*-[Δ*1*] allele with the *his3*Δ*1* allele present in strain BY4741 derivatives does not result in a functional *HIS3* allele [36], Strains containing Mediator subunit gene deletions were constructed via lithium acetate transformation with a *KanMX* allele as described [86, 87].

A chromosomal Ty1*kanMXAI* element was introduced into strain BY4741 by inducing the expression of a Ty1*kanMXAI* element fused to the *GAL1* promoter on a yeast URA3-marked 2μ vector (plasmid pGTy1*kanMXAI* [39]), as described previously [19]. Briefly, a transformant of strain BY4741 harboring plasmid pGTy1*kanMXAI* was grown on SC-URA 2% glucose 2% raffinose agar at 20°C. Twenty single colonies were picked and struck for single colonies on SC-URA 2% glucose and grown at 30°C. A single Ura+ colony harboring cells had maintained the pGTy1*kanMXAI* plasmid throughout galactose-induction was picked from each streak and struck for single colonies on YPD and grown at 30°C. A single colony was picked from each YPD streak and used to make a 1 cm^2^ patch on YPD at 30°C. Each patch was replicated to a fresh YPD agar and grown for three days at 20°C, and then replicated to YPD agar containing 200 μg/ml G418 and grown at 30°C for three days. Strains harboring a chromosomal Ty1*kanMXAI* element were identified by the appearance of G418^R^ papillae. A strain with a representative number of G418^R^ papillae, JC6464, was chosen for further analysis. A *med15Δ:URA3* derivative of strain JC6464 was constructed by PCR-mediated gene disruption.

Plasmid Ycp50-Ty1*his3AI*-[Δ1] is a *URA3-CEN*plasmid containing a Ty1 *his3AI-[Δ1]* element, constructed by replacing the *ClaI* fragment containing the *his3AI* allele in pBDG633 with a *ClaI* fragment containing *his3AI-[Δ1]* and kindly provided by Dr. David Garfinkel [88]. Plasmid pBJC1250 is a *LEU2-CEN*vector containing a *Ty1*his3AI*-[Δ1]* element wherein the U3 region of the 5′ LTR was replaced with a *TEF1* promoter (herein referred to as P_*TEF1*_-Ty1*his3AI*; see Fig 5). Plasmid pBJC80, herein referred to as pGTy1ΔPOL, has been described previously [69].

### Transposition Frequency Assay

Ty1 retromobility was determined as previously described [19]. Individual colonies from strains were grown in triplicate in YPD broth at 30°C overnight. Each culture was then diluted in quadruplicate by a factor of 1000 in YPD broth and grown at 20°C to an optical density beyond log growth phase. 1μL of a 1:1000 dilution of each of the resulting 12 cultures was plated on YPD agar to provide an accurate representation of the cell density. In parallel, 100μL to 1mL of each culture was plated on SC-HIS agar. All plates were grown at 30°C for 3-4 days. Mobility frequency was calculated as a ratio of the number of His+ colony forming units to the number of colony forming units in each culture as represented by the number of colonies growing on YPD agar. For strains for which no His+ prototrophs were observed, mobility was reported as an upper limit equal to the ratio of (1/the total number of colony forming units in all three biological replicates).

For strains containing a plasmid Ty1*his3AI* element, the above protocol was modified such that cultures were grown in their respective selective media (SC-URA or SC-LEU with 2% glucose) at 30°C until confluent, diluted 1:1000 and grown at 20°C in YPD until confluent, and plated on their respective dropout media (SC-URA or SC-LEU with 2% glucose) as well as the corresponding media lacking histidine (SC-URA-HIS or SC-LEU-HIS with 2% glucose).

For strains containing a chromosomal Ty1*kanMXAI* element and plasmid pBJC1250 containing the P_*TEF1*_-Ty1*his3AI* element, the above protocol was modified such that cultures were grown in SC-LEU 2% glucose at 30°C overnight. Cultures were diluted 1: 1000 into YPD broth and separated into 12 independent cultures that were grown for three days at 20°C. A 1:1000 dilution of each culture was plated on YPD agar to determine the number of colony forming units in each culture, and aliquots of each culture were plated YPD agar containing 200 μg/ml G418 to determine the number of G418^R^ colony forming units per culture, and SC-HIS 2% glucose to determine the number of His+ colony forming units per culture.

### RNA Purification

Cells were grown in YPD broth at 20°C to mid-log phase. Total cellular RNA was extracted using a hot phenol/chloroform extraction protocol [89]. PolyA^+^ RNA was purified from 250μg to 1mg of total cellular RNA using the Magnetic mRNA Isolation Kit (New England Biolabs) following the manufacturer’s protocol.

### Northern Blot

A 20μg aliquot of total cellular RNA was separated on a 1% Seakem GTG agarose gel as previously described [90]. Three to six μg of poly A+ RNA was separated on an 0.8% Seakem GTG agarose gel as previously described [90]. Separated RNA was transferred to a Hybond XL membrane (GE Healthcare) using a gradient of 6X SSC to 10X SSC overnight at room temperature. Synthesis of ^32^P-labeled RNA riboprobes was carried out in vitro using SP6 or T7 polymerase (New England Biolabs). Membranes were incubated with probes in NorthernMax PreHyb Buffer (Ambion) at 65 °C overnight. Images were scanned using a Typhoon 9400 scanner, and quantified using ImageQuant software (Molecular Dynamics, Sunnyvale, CA).

### Western Blot

Cultures were grown at 20°C in YPD for one cell doubling (OD_600_ 0.3 to OD_600_ 0.6) after dilution from overnight cultures grown at 30°C. Protein was extracted from total cell lysates as previously described [91] and resolved on a 10% SDS-PAGE gel. When resolving p18- and p22-Gag, a 15% SDS-PAGE gel was used. Protein was then transferred to a polyvinylidene difluoride (PVDF) membrane. Membranes were blocked in a 5% nonfat milk solution dissolved in phosphate buffered saline (PBS) with 0.1% TWEEN 20. Membranes were then incubated in 0.5% nonfat milk in PBS with 0. 1% TWEEN 20 with a 1:7500 dilution of affinity-purified anti-VLP antisera [53] to detect Gag, a 1:5000 dilution of a polyclonal antibody specific to p18-Gag (a gift from David Garfinkel, described in [24]), a 1:7500 dilution of anti GAPDH monoclonal antibody (Thermo Fisher Scientific), or a 1:5000 dilution of anti actin monoclonal antibody (Abcam). Med17-myc was detected using a 1:1000 dilution of a monoclonal antibody to c-myc (Sigma-Aldrich). Membranes were subsequently incubated with horseradish peroxidase (HRP)-conjugated secondary antibodies (Millipore). Following terminal washes, membranes were incubated with SuperSignal West Pico chemiluminescence substrate (Pierce, Thermo Fisher Scientific), and exposed to film (Kodak). Antibody was stripped from membranes as described previously [92]. Images were developed on film using a Model SRX-101A Medical Scanner (Konica Minolta) and scanned using a Cannon MP480 scanner. Protein bands were quantified using ImageJ (NIH). Quantification was performed using film exposed for different durations to ensure that measurements were done within the linear response range.

### Southern Blot

Cultures were grown past log growth phase at 20°C in YPD broth. Total genomic DNA was isolated as previously described [15, 93], and digested with *Sph I* endonuclease. Digested genomic DNA was then fractionated by gel electrophoresis on a 1% GTG agarose gel and subjected to Southern blot analysis with a ^32^P-labeled riboprobe specific for *POL* as described previously [15, 37].

### cDNA Synthesis and Estimation of pGTy1ΔPOL expression

Polyadenylated RNA was used to synthesize cDNA using the First Strand cDNA Synthesis Kit (Affymetrix). 100ng of polyA+ mRNA was used per reaction, as were 0.5μM concentration each of primers specific to the *ΔPOL* region of pGTy1ΔPOL (5′-CCACCCATAATGTAATAGATCTATCGATTCTAGAC-3′) and to *ACT1* (5′-ATCGTCCCAGTTGGTGACAATACC-3′). Reactions were performed according to manufacturer protocols, and run at 44°C for 1 hour, followed by incubation at 92°C for 10 minutes. For comparison of levels of full-length Ty1 and Ty1i transcripts generated from pGTy1ΔPOL (Fig 6), cDNA was PCR amplified using primers binding 250 bp upstream of the Ty1i TSS (5′-GATTCATCCTCAGCGGACTCTG-3′), or 250 bp downstream of Ty1i TSS (5′-AGAAGAATGATTCTCGCAGC-3′), and the *ΔPOL* region of the element (5′-CCACCCATAATGTAATAGATCTATCGATTCTAGAC-3′). PCR product was isolated at different cycle numbers and compared with amplification of *ACT1* (forward primer: 5′-GGTTCTGGTATGTGTAAAGCCGGT-3′; reverse primer: 5′-ATCGTCCCAGTTGGTGACAATACC-3′) to control for relative cDNA abundance. A control set of PCR amplifications was performed using genomic DNA prepared from the same strain.

Expression of pGTy1ΔPOL relative to expression of *ACT1* was estimated as follows: Amplification of *ACT1* cDNA was observed at 5-6 fewer cycles than the average amplification for Ty1 and Ty1i, whereas amplification of gDNA was about equal to this average (Fig. 6B), thus indicating conservatively a 32-fold greater amount of *ACT1* transcript. The amplicon for *ACT1* is about 200 bp, while the amplicons for Ty1ΔPOL and Ty1iΔPOL are each about 3 kb, and so an equal intensity for *ACT1* and Ty1ΔPOL corresponds to about a 15-fold greater molar quantity of *ACT1.* Combining these ratios indicates that *ACT1* transcript levels are approximately 450 times greater than those for Ty1 POL.

### ChIP-seq

For analysis of Mediator occupancy at Ty1 elements (Fig 8), chromatin immunoprecipitation followed by high throughput sequencing (ChIP-seq) was performed using strains in which Mediator subunits Med15 or Med17 carried 13-myc epitope tags and which were engineered to allow Kin28 inactivation by the anchor away technique (S1 Table) [63]. For anchor away experiments, yeast were grown in YPD to an OD_600_ of 0.8. Rapamycin was then added to a concentration of 1 μg/mL (from a 1 mg/mL stock in ethanol stored at -20°C for not more than one month) and cultures allowed to grow one hr at 30°C prior to cross-linking. ChIP against epitope-tagged Mediator subunits was carried out as described previously [61], using 2 μg of anti-myc antibody (clone 9E10, Sigma) and protein G sepharose beads for capture (GE Healthcare). Library preparation for Illumina sequencing was performed using the NEBNext Ultra II library preparation kit (New England Biolabs) according to manufacturer’s directions. Libraries were bar-coded using NEXTflex barcodes (BIOO Scientific) and sequenced on the Illumina NextSeq platform at the Wadsworth Center, New York State Department of Health.

Unfiltered paired-end sequencing reads were aligned to the *S. cerevisiae* reference genome (Saccer3) by using BWA [94]. Up to one mismatch was allowed for each aligned read; reads mapping to multiple locations were retained and randomly assigned. Because full-length Ty1 elements share sequences with 266 Ty1 delta elements (263 of which are <350 bp in length), some reads from the first ~350 bp of Ty1 elements will also be mapped to these elements. Duplicate reads were removed based on paired end information. Occupancy profiles for Ty1 elements were generated by averaging the signal of all 31 Ty1 elements; thus, behavior of individual elements cannot be assessed. The occupancy is plotted on the window from 2kb upstream of TSS to 2kb downstream of TES (Fig 5 and 6). In each Ty1 element ORF, from TSS+1kb to TES-1kb, the region is divided into 100 bins and the average occupancy of each bin was calculated. For the flanking regions, the occupancy was calculated for each base pair.

### Data Deposition

ChIP-seq data has been deposited in Arrayexpress under accession number E-MTAB-5824. Data used in Figure 4B is available from the NCBI Sequence Read Archive under accession number SRP047524.

## Acknowledgements

We thank Todd Benziger for help with strain construction, and David Garfinkel, PhD (University of Georgia, Athens, Georgia, USA) for the donation of the anti-p18 antibody and helpful advice on its use. Additionally, we thank the Wadsworth Center Media and Tissue Culture Core and Applied Genomic Technologies Core facilities.

## S1 Fig. Mediator subunit deletions have only modest effects on levels of p49/p45 Gag protein levels

(A) Western blot of total cell lysates from wild-type, *spt3Δ*, and Mediator subunit deletion strains probed for Gag using an anti-VLP antibody. (B) Quantitation of p49/45 levels relative to α-actin from (A), normalized to WT.

## S2 Fig. Northern analysis of *Ty1*his3AI** expressed from the LTR or *TEF1* promoter

The blot was probed with a sense-strand *HIS3* riboprobe to detect Ty1*his3AI* and Ty1i*his3AI*RNA. All lanes shown are from a single gel. Note the absence of any band below the Ty1*his3AI* transcript (compare Fig. 5A). The values reported in the graph are the average ratio of Ty1 *his3AI* RNA relative to *PYK1* RNA in two biological replicates. Bars are color-coded as in Fig. 2A: gray, WT and *spt3A*; red, Mediator head module deletion; blue, middle module deletion; yellow, tail module deletion. All error bars represent s.d.

## S3 Fig. Occupancy of Mediator over Ty1 elements determined by ChIP-seq

Top, untagged control subjected to ChIP-seq using anti-myc antibody. Bottom, occupancy of myc-tagged Med15 and Med17 over averaged Ty1 elements in *Med18Δ kin28-AA* yeast. Ty1 elements begin at 0 kb on the x-axis, and the Ty1 TSS at +238 and Ty1i TSS at +1000 are marked by green bars on the x-axis. Note the absence of any Mediator peak at the Ty1i TSS.

## S4 Fig. Occupancy of Pol II and Med17 over Ty1 elements in wild type and tail module deletion mutants

Data is the same as in Fig. 5 and 6.

## S5 Fig. Quantitation of Med17-myc levels in wild-type and Mediator mutant yeast strains

(A) Western blot of total cell lysates from wild-type and mutant yeast expressing c-Myc tagged Med17 (first four lanes) and from an untagged control strain (last lane), probed using an antibody against c-Myc and against GAPDH. (B) Quantitation of two biological replicate experiments as in (A).

## S6 Fig. Quantitation of Ty1ΔPOL and Ty1iΔPOL transcripts expressed from pGTy1ΔPOL in yeast grown in glucose medium

Top: Schematic of the GAL1:Ty1ΔPOL cassette in pGTy1ΔPOL showing forward primer locations for detection of Ty1 RNA (blue) versus Ty1i RNA (red). Note that because Ty1i is contained within Ty1, the Ty1i primer reports both Ty1i and Ty1 transcripts. Both amplifications utilized the same reverse primer (purple), that crosses the deletion junction and contains sequences unique to the pGTy1ΔPOL element. A reverse primer specific for *ACT1* was also used to synthesize cDNA used as a template for the PCR amplification. No PCR product was detected using the ΔPOL reverse primer when RNA from yeast lacking pGTy1ΔPOL was used as template. Bottom: Reverse Transciption-PCR reactions using polyA^+^ RNA isolated from strains of the indicated genotype bearing plasmid pGTy1ΔPOL, and grown in glucose-containing broth. Aliquots were taken from reactions at the indicated number of cycles and analyzed by agarose gel electrophoresis. RT-PCR amplification products using Ty1, Ty1i, and *ACT1* primers are indicated. We do not know the origin of the apparently spurious, lower molecular weight bands observed. The same WT samples were used for all panels; results were similar for a second biological replicate of all three samples (WT, *Med20Δ*, and *med3*).

**S1 Table.** Yeast strains used in this work.

## References

1. Curcio MJ, Derbyshire KM. The outs and ins of transposition: from mu to kangaroo. Nat Rev Mol Cell Biol. 2003;4(11):865–77.

2. Biemont C. A brief history of the status of transposable elements: from junk DNA to major players in evolution. Genetics. 2010;186(4):1085–93.

3. Gogvadze E, Buzdin A. Retroelements and their impact on genome evolution and functioning. Cell Mol Life Sci. 2009;66(23):3727–42.

4. Ivancevic AM, Kortschak RD, Bertozzi T, Adelson DL. LINEs between Species: Evolutionary Dynamics of LINE-1 Retrotransposons across the Eukaryotic Tree of Life. Genome Biol Evol. 2016;8(11):3301–22.

5. Kazazian HH, Jr. Mobile elements: drivers of genome evolution. Science. 2004;303(5664):1626–32.

6. Hancks DC, Kazazian HH, Jr. Roles for retrotransposon insertions in human disease. Mob DNA. 2016;7:9.

7. Papasotiriou I, Pantopikou K, Apostolou P. L1 retrotransposon expression in circulating tumor cells. PLoS One. 2017;12(2):e0171466.

8. Scott EC, Gardner EJ, Masood A, Chuang NT, Vertino PM, Devine SE. A hot LI retrotransposon evades somatic repression and initiates human colorectal cancer. Genome Res. 2016;26(6):745–55.

9. Malik HS, Henikoff S, Eickbush TH. Poised for contagion: evolutionary origins of the infectious abilities of invertebrate retroviruses. Genome Res. 2000;10(9):1307–18.

10. Irwin B, Aye M, Baldi P, Beliakova-Bethell N, Cheng H, Dou Y, et al Retroviruses and yeast retrotransposons use overlapping sets of host genes. Genome Res. 2005;15(5):641–54.

11. Dutko JA, Schafer A, Kenny AE, Cullen BR, Curcio MJ. Inhibition of a yeast LTR retrotransposon by human APOBEC3 cytidine deaminases. Curr Biol. 2005;15(7):661–6.

12. Maxwell PH, Curcio MJ. Host factors that control long terminal repeat retrotransposons in Saccharomyces cerevisiae: implications for regulation of mammalian retroviruses. Eukaryot Cell. 2007;6(7):1069–80.

13. Brass AL, Dykxhoorn DM, Benita Y, Yan N, Engelman A, Xavier RJ, et al Identification of host proteins required for HIV infection through a functional genomic screen. Science. 2008;319(5865):921–6.

14. Zhou H, Xu M, Huang Q, Gates AT, Zhang XD, Castle JC, et al Genome-scale RNAi screen for host factors required for HIV replication. Cell Host Microbe. 2008;4(5):495–504.

15. Dutko JA, Kenny AE, Gamache ER, Curcio MJ. 5’ to 3’ mRNA decay factors colocalize with Ty1 gag and human APOBEC3G and promote Ty1 retrotransposition. J Virol. 2010;84(10):5052–66.

16. Checkley MA, Mitchell JA, Eizenstat LD, Lockett SJ, Garfinkel DJ. Ty1 gag enhances the stability and nuclear export of Ty1 mRNA. Traffic. 2013;14(1):57–69.

17. Okada A, Iwatani Y. APOBEC3G-Mediated G-to-A Hypermutation of the HIV-1 Genome: The Missing Link in Antiviral Molecular Mechanisms. Front Microbiol. 2016;7:2027.

18. Curcio MJ, Lutz S, Lesage P. The Ty1 LTR-retrotransposon of budding yeast. Microbiol Spectr. 2015;3(2):1–35.

19. Curcio MJ, Garfinkel DJ. Single-step selection for Ty1 element retrotransposition. Proc Natl Acad Sci USA. 1991;88(3):936–40.

20. Basehoar AD, Zanton SJ, Pugh BF. Identification and distinct regulation of yeast TATA box-containing genes. Cell. 2004;116(5):699–709.

21. Kenna MA, Brachmann CB, Devine SE, Boeke JD. Invading the yeast nucleus: a nuclear localization signal at the C terminus of Ty1 integrase is required for transposition in vivo. Mol Cell Biol. 1998;18(2):1115–24.

22. Moore SP, Rinckel LA, Garfinkel DJ. A Ty1 integrase nuclear localization signal required for retrotransposition. Mol Cell Biol. 1998;18(2):1105–14.

23. Happel AM, Swanson MS, Winston F. The SNF2, SNF5 and SNF6 genes are required for Ty transcription in Saccharomyces cerevisiae. Genetics. 1991;128(1):69–77.

24. Saha A, Mitchell JA, Nishida Y, Hildreth JE, Ariberre JA, Gilbert WV, et al A trans-dominant form of Gag restricts Ty1 retrotransposition and mediates copy number control. J Virol. 2015;89(7):3922–38.

25. Winston F, Durbin KJ, Fink GR. The SPT3 gene is required for normal transcription of Ty elements in S. cerevisiae. Cell. 1984;39(3 Pt 2):675–82.

26. Garfinkel DJ, Nyswaner K, Wang J, Cho JY. Post-transcriptional cosuppression of Ty1 retrotransposition. Genetics. 2003;165(1):83–99.

27. Matsuda E, Garfinkel DJ. Posttranslational interference of Ty1 retrotransposition by antisense RNAs. Proc Natl Acad Sci USA. 2009;106(37):15657–62.

28. Nishida Y, Pachulska-Wieczorek K, Blaszczyk L, Saha A, Gumna J, Garfinkel DJ, et al Ty1 retroviruslike element Gag contains overlapping restriction factor and nucleic acid chaperone functions. Nucleic Acids Res. 2015;43(15):7414–31.

29. Tucker JM, Larango ME, Wachsmuth LP, Kannan N, Garfinkel DJ. The Ty1 Retrotransposon Restriction Factor p22 Targets Gag. Plos Genet. 2015;11(10):el005571.

30. Tucker JM, Garfinkel DJ. Ty1 escapes restriction by the self-encoded factor p22 through mutations in capsid. Mob Genet Elements. 2016;6(2):e1154639.

31. Berretta J, Pinskaya M, Morillon A. A cryptic unstable transcript mediates transcriptional trans-silencing of the Ty1 retrotransposon in S. cerevisiae. Genes Dev. 2008;22(5):615–26.

32. Suresh S, Ahn HW, Joshi K, Dakshinamurthy A, Kananganat A, Garfinkel DJ, et al Ribosomal protein and biogenesis factors affect multiple steps during movement of the Saccharomyces cerevisiae Ty1 retrotransposon. Mob DNA. 2015;6:22.

33. Garfinkel DJ, Tucker JM, Saha A, Nishida Y, Pachulska-Wieczorek K, Blaszczyk L, et al A self-encoded capsid derivative restricts Ty1 retrotransposition in Saccharomyces. Curr Genet. 2016;62(2):321–9.

34. Dakshinamurthy A, Nyswaner KM, Farabaugh PJ, Garfinkel DJ. BUD22 affects Ty1 retrotransposition and ribosome biogenesis in Saccharomyces cerevisiae. Genetics. 2010;185(4):1193–205.

35. Malagon F, Jensen TH. The T body, a new cytoplasmic RNA granule in Saccharomyces cerevisiae. Mol Cell Biol. 2008;28(19):6022–32.

36. Nyswaner KM, Checkley MA, Yi M, Stephens RM, Garfinkel DJ. Chromatin-associated genes protect the yeast genome from Ty1 insertional mutagenesis. Genetics. 2008;178(1):197–214.

37. Scholes DT, Banerjee M, Bowen B, Curcio MJ. Multiple regulators of Ty1 transposition in Saccharomyces cerevisiae have conserved roles in genome maintenance. Genetics. 2001;159(4):1449–65.

38. Griffith JL, Coleman LE, Raymond AS, Goodson SG, Pittard WS, Tsui C, et al Functional genomics reveals relationships between the retrovirus-like Ty1 element and its host Saccharomyces cerevisiae. Genetics. 2003;164(3):867–79.

39. Curcio MJ, Kenny AE, Moore S, Garfinkel DJ, Weintraub M, Gamache ER, et al S-phase checkpoint pathways stimulate the mobility of the retrovirus-like transposon Ty1. Mol Cell Biol. 2007;27(24):8874–85.

40. Allen BL, Taatjes DJ. The Mediator complex: a central integrator of transcription. Nat Rev Mol Cell Biol. 2015;16(3):155–66.

41. Ansari SA, He Q, Morse RH. Mediator complex association with constitutively transcribed genes in yeast. P Natl Acad Sci USA. 2009;106(39):16734–9.

42. Ansari SA, Morse RH. Mechanisms of Mediator complex action in transcriptional activation. Cell Mol Life Sci. 2013;70(15):2743–56.

43. Esnault C, Ghavi-Helm Y, Brun S, Soutourina J, Van Berkum N, Boschiero C, et al Mediator-dependent recruitment of TFIIH modules in preinitiation complex. Mol Cell. 2008;31(3):337–46.

44. Grunberg S, Henikoff S, Hahn S, Zentner GE. Mediator binding to UASs is broadly uncoupled from transcription and cooperative with TFIID recruitment to promoters. EMBO J. 2016;35(22):2435–46.

45. Jeronimo C, Langelier MF, Bataille AR, Pascal JM, Pugh BF, Robert F. Tail and Kinase Modules Differently Regulate Core Mediator Recruitment and Function In Vivo. Mol Cell. 2016;64(3):455–66.

46. Kornberg RD. Mediator and the mechanism of transcriptional activation. Trends Biochem Sci. 2005;30(5):235–9.

47. Robinson PJ, Trnka MJ, Bushnell DA, Davis RE, Mattei PJ, Burlingame AL, et al Structure of a Complete Mediator-RNA Polymerase II Pre-Initiation Complex. Cell. 2016;166(6):1411–22 el6.

48. Plaschka C, Nozawa K, Cramer P. Mediator Architecture and RNA Polymerase II Interaction. J Mol Biol. 2016;428(12):2569–74.

49. Robinson PJ, Trnka MJ, Pellarin R, Greenberg CH, Bushnell DA, Davis R, et al Molecular architecture of the yeast Mediator complex. Elife. 2015;4.

50. Tsai KL, Tomomori-Sato C, Sato S, Conaway RC, Conaway JW, Asturias FJ. Subunit architecture and functional modular rearrangements of the transcriptional mediator complex. Cell. 2014;157(6):1430–44.

51. Wang X, Sun Q, Ding Z, Ji J, Wang J, Kong X, et al Redefining the modular organization of the core Mediator complex. Cell Res. 2014;24(7):796–808.

52. Jeronimo C, Robert F. The Mediator Complex: At the Nexus of RNA Polymerase II Transcription. Trends Cell Biol. 2017;27(10):765–83.

53. Risler JK, Kenny AE, Palumbo RJ, Gamache ER, Curcio MJ. Host co-factors of the retrovirus-like transposon Ty1. Mob DNA. 2012;3(1):12.

54. Boeke JD, Styles CA, Fink GR. Saccharomyces cerevisiae SPT3 gene is required for transposition and transpositional recombination of chromosomal Ty elements. Mol Cell Biol. 1986;6(11):3575–81.

55. Myers LC, Gustafsson CM, Hayashibara KC, Brown PO, Kornberg RD. Mediator protein mutations that selectively abolish activated transcription. Proc Natl Acad Sci USA. 1999;96(1):67–72.

56. Zhang F, Sumibcay L, Hinnebusch AG, Swanson MJ. A triad of subunits from the Galll/tail domain of Srb mediator is an in vivo target of transcriptional activator Gcn4p. Mol Cell Biol. 2004;24(15):6871–86.

57. Ansari SA, Ganapathi M, Benschop JJ, Holstege FC, Wade JT, Morse RH. Distinct role of Mediator tail module in regulation of SAGA-dependent, TATA-containing genes in yeast. EMBO J. 2012;31(1):44–57.

58. van de Peppel J, Kettelarij N, van Bakel H, Kockelkorn TT, van Leenen D, Holstege FC. Mediator expression profiling epistasis reveals a signal transduction pathway with antagonistic submodules and highly specific downstream targets. Mol Cell. 2005;19(4):511–22.

59. Ansari SA, Morse RH. Selective role of Mediator tail module in the transcription of highly regulated genes in yeast. Transcription. 2012;3:110–4.

60. Kemmeren P, Sameith K, van de Pasch LA, Benschop JJ, Lenstra TL, Margaritis T, et al Large-scale genetic perturbations reveal regulatory networks and an abundance of gene-specific repressors. Cell. 2014;157(3):740–52.

61. Paul E, Zhu ZI, Landsman D, Morse RH. Genome-wide association of mediator and RNA polymerase II in wild-type and mediator mutant yeast. Mol Cell Biol. 2015;35(1):331–42.

62. Kadosh D, Struhl K. Targeted recruitment of the Sin3-Rpd3 histone deacetylase complex generates a highly localized domain of repressed chromatin in vivo. Mol Cell Biol. 1998;18(9):5121–7.

63. Haruki H, Nishikawa J, Laemmli UK. The anchor-away technique: rapid, conditional establishment of yeast mutant phenotypes. Mol Cell. 2008;31(6):925–32.

64. Wong KH, Jin Y, Struhl K. TFIIH phosphorylation of the Pol II CTD stimulates mediator dissociation from the preinitiation complex and promoter escape. Mol Cell. 2014;54(4):601–12.

65. Jeronimo C, Robert F. Kin28 regulates the transient association of Mediator with core promoters. Nat Struct Mol Biol. 2014;21(5):449–55.

66. Dang Y, Cheng J, Sun X, Zhou Z, Liu Y. Antisense transcription licenses nascent transcripts to mediate transcriptional gene silencing. Genes Dev. 2016;30(21):2417–32.

67. Martens JA, Laprade L, Winston F. Intergenic transcription is required to repress the Saccharomyces cerevisiae SER3 gene. Nature. 2004;429(6991):571–4.

68. Prescott EM, Proudfoot NJ. Transcriptional collision between convergent genes in budding yeast. Proc Natl Acad Sci USA. 2002;99(13):8796–801.

69. Curcio MJ, Garfinkel DJ. Heterogeneous functional Ty1 elements are abundant in the Saccharomyces cerevisiae genome. Genetics. 1994;136(4):1245–59.

70. Huisinga KL, Pugh BF. A genome-wide housekeeping role for TFIID and a highly regulated stress-related role for SAGA in Saccharomyces cerevisiae. Mol Cell. 2004;13(4):573–85.

71. Munchel SE, Shultzaberger RK, Takizawa N, Weis K. Dynamic profiling of mRNA turnover reveals gene-specific and system-wide regulation of mRNA decay. Mol Biol Cell. 2011;22(15):2787–95.

72. Huang Q, Purzycka KJ, Lusvarghi S, Li D, Legrice SF, Boeke JD. Retrotransposon Ty1 RNA contains a 5’-terminal long-range pseudoknot required for efficient reverse transcription. RNA. 2013;19(3):320–32.

73. Yu G, Fassler JS. SPT13 (GAL11) of Saccharomyces cerevisiae negatively regulates activity of the MCM1 transcription factor in Ty1 elements. Mol Cell Biol. 1993;13(1):63–71.

74. Sokolowski M, Chynces M, deHaro D, Christian CM, Belancio VP. Truncated 0RF1 proteins can suppress LINE-1 retrotransposition in trans. Nucleic Acids Res. 2017;45(9):5294–308.

75. Persson J, Steglich B, Smialowska A, Boyd M, Bornholdt J, Andersson R, et al Regulating retrotransposon activity through the use of alternative transcription start sites. EMBO Rep. 2016;17(5):753–68.

76. Miller C, Matic I, Maier K, Schwalb B, Roether S, Straesser K, et al Mediator phosphorylation prevents stress response transcription during non-stress conditions. J Biol Chem. 2012.

77. Morillon A, Benard L, Springer M, Lesage P. Differential effects of chromatin and Gcn4 on the 50-fold range of expression among individual yeast Ty1 retrotransposons. Mol Cell Biol. 2002;22(7):2078–88.

78. Servant G, Pennetier C, Lesage P. Remodeling yeast gene transcription by activating the Ty1 long terminal repeat retrotransposon under severe adenine deficiency. Mol Cell Biol. 2008;28(17):5543–54.

79. Sacerdot C, Mercier G, Todeschini AL, Dutreix M, Springer M, Lesage P. Impact of ionizing radiation on the life cycle of Saccharomyces cerevisiae Ty1 retrotransposon. Yeast. 2005;22(6):441–55.

80. Staleva Staleva L, Venkov P. Activation of Ty transposition by mutagens. Mutat Res. 2001;474(1-2):93–103.

81. Todeschini AL, Morillon A, Springer M, Lesage P. Severe adenine starvation activates Ty1 transcription and retrotransposition in Saccharomyces cerevisiae. Mol Cell Biol. 2005;25(17):7459–72.

82. Xu H, Boeke JD. Inhibition of Ty1 transposition by mating pheromones in Saccharomyces cerevisiae. Mol Cell Biol. 1991;11(5):2736–43.

83. Checkley MA, Nagashima K, Lockett SJ, Nyswaner KM, Garfinkel DJ. P-body components are required for Ty1 retrotransposition during assembly of retrotransposition-competent virus-like particles. Mol Cell Biol. 2010;30(2):382–98.

84. Drinnenberg IA, Weinberg DE, Xie KT, Mower JP, Wolfe KH, Fink GR, et al RNAi in budding yeast. Science. 2009;326(5952):544–50.

85. Mou Z, Kenny AE, Curcio MJ. Hos2 and Set3 promote integration of Ty1 retrotransposons at tRNA genes in Saccharomyces cerevisiae. Genetics. 2006;172(4):2157–67.

86. Brachmann CB, Davies A, Cost GJ, Caputo E, Li J, Hieter P, et al Designer deletion strains derived from Saccharomyces cerevisiae S288C: a useful set of strains and plasmids for PCR-mediated gene disruption and other applications. Yeast. 1998;14(2):115–32.

87. Winzeler EA, Shoemaker DD, Astromoff A, Liang H, Anderson K, Andre B, et al Functional characterization of the S. cerevisiae genome by gene deletion and parallel analysis. Science. 1999;285(5429):901–6.

88. Lee BS, Lichtenstein CP, Faiola B, Rinckel LA, Wysock W, Curcio MJ, et al Posttranslational inhibition of Ty1 retrotransposition by nucleotide excision repair/transcription factor TFIIH subunits Ssl2p and Rad3p. Genetics. 1998;148(4):1743–61.

89. Schmitt ME, Brown TA, Trumpower BL. A rapid and simple method for preparation of RNA from Saccharomyces cerevisiae. Nucleic Acids Res. 1990;18(10):3091–2.

90. Maxwell PH, Coombes C, Kenny AE, Lawler JF, Boeke JD, Curcio MJ. Ty1 mobilizes subtelomeric Y’ elements in telomerase-negative Saccharomyces cerevisiae survivors. Mol Cell Biol. 2004;24(22):9887–98.

91. Kushnirov W. Rapid and reliable protein extraction from yeast. Yeast. 2000;16(9):857–60.

92. Doh JH, Lutz S, Curcio MJ. Co-translational localization of an LTR-retrotransposon RNA to the endoplasmic reticulum nucleates virus-like particle assembly sites. Plos Genet. 2014;10(3):e1004219.

93. Conte D, Jr., Barber E, Banerjee M, Garfinkel DJ, Curcio MJ. Posttranslational regulation of Ty1 retrotransposition by mitogen-activated protein kinase Fus3. Mol Cell Biol. 1998;18(5):2502–13.

94. Seoighe C, Wolfe KH. Updated map of duplicated regions in the yeast genome. Gene. 1999;238(1):253–61.

